# From Plant Detection to Satellite Mapping: A Multi-Scale AI Toolkit for Ragweed Surveillance under Climate Change

**DOI:** 10.64898/2026.09.28.755009

**Authors:** Lorenzo F. León Gutiérrez, Cristofer Ramírez, Alejandra Henríquez, Sebastián Contreras

## Abstract

Climate change is reshaping weed population dynamics, creating non-stationary management challenges for which static decision rules are insufficient. This chapter presents and evaluates an open surveillance toolkit of artificial intelligence tools for recognizing and mapping Ambrosia artemisiifolia (common ragweed) at complementary spatial scales, tested across two phenological phases (seedling, September 2024; adult, December 2024) in a lentil paddock in central Chile. At the field scale, a YOLOv11 detection model coupled with Slicing Aided Hyper Inference achieved mAP50 of 0.886. Cross-domain experiments revealed that single-site models collapse catastrophically when deployed in new geographies (mAP50 dropping to 0.108), but multi-domain training recovers performance to 0.874. At the satellite scale, PRESTO foundation model embeddings correlated strongly with ground-truth weed density (r = 0.739); geographically weighted regression explained up to 91% of local density variance. All methods are released as the ragweed-ai-toolkit (https://github.com/agroia-lab/ragweed-ai-toolkit), a modular open-source Python package enabling adaptation to new species and geographies.

## 1 Introduction

Climate change is reshaping the spatial and temporal dynamics of weed populations across agricultural landscapes worldwide. Rising atmospheric CO_2_ concentrations, shifting temperature regimes, and altered precipitation patterns are enabling range expansions of invasive species into previously uncolonized regions while modifying the phenology, growth rates, and competitive interactions of resident weed communities (Lake et al., 2017). Common ragweed (*Ambrosia artemisiifolia* L.) exemplifies these dynamics: projections indicate that ragweed pollen sensitization in Europe will more than double by 2050 under moderate climate scenarios, with airborne pollen concentrations projected to increase fourfold through combined range expansion and climate-driven phenological shifts (Hamaoui-Laguel et al., 2015), driving annual healthcare costs into billions of euros across the continent (Lake et al., 2017; Schaffner et al., 2020). Analysis of North American pollen records confirms that pollen seasons have already lengthened by approximately 20 days and airborne concentrations have increased by 21% since 1990, with anthropogenic climate change responsible for roughly half of the trend (Anderegg et al., 2021). Under continued warming, pollen emissions are projected to increase by 16–40% from climate effects alone and by up to 200% when CO_2_ fertilization is included (Zhang & Steiner, 2022). Experimental evidence demonstrates that elevated urban CO_2_ concentrations amplify ragweed above-ground biomass by up to 189% (Ziska et al., 2003), while agricultural source populations feed urban pollen loads through wind dispersal over hundreds of kilometers (Essl et al., 2015). More broadly, long-term field data from the Broadbalk experiment demonstrate that the competitive advantage of weeds over crops has increased steadily since 1969 in response to warming and shorter cultivar stature (Storkey et al., 2021), and weeds as a functional group adapt faster than crops to changing conditions owing to greater phenotypic plasticity (Anwar et al., 2021). These trends create a dual management imperative that links in-field weed control to public health outcomes—a linkage that conventional, static decision frameworks are ill-equipped to address under non-stationary environmental conditions.

Effective management of ragweed infestations depends fundamentally on surveillance: knowing *where* populations occur at sufficient spatial and temporal resolution to intervene before reproductive maturity and peak pollen dispersal. Manual scouting— walking fields to identify and map problem patches—remains the practical bottleneck: it is too slow to cover landscape-scale areas before the flowering window, too expensive for routine pre-season monitoring, and unreliable across heterogeneous field conditions. In Chile, where *A. artemisiifolia* is confirmed as introduced and where ecological niche expansion exceeds that observed on any other continent (Song et al., 2023), this surveillance deficit translates directly into undetected infestations that reach seed production unchecked. Automating weed surveillance at multiple spatial scales—from individual plant detection to satellite-derived density maps—is therefore a prerequisite for the source-reduction strategies described above.

Artificial intelligence tools capable of addressing this surveillance gap have advanced rapidly in the past decade, yet they remain largely compartmentalized. Object detection architectures from the You Only Look Once (YOLO) family have demonstrated reliable weed identification in diverse cropping systems (Rai et al., 2023; Hu et al., 2024), while geostatistical methods enable mapping of continuous weed density surfaces from point observations. At the satellite scale, vegetation indices derived from Sentinel-2 imagery offer coarse spatial monitoring, and foundation models pre-trained on remote sensing time series—notably the Pretrained Remote Sensing Transformer (PRESTO) (Tseng et al., 2023)—have begun to demonstrate general-purpose representation learning for agricultural applications. Despite these individual advances, no integrated surveillance framework links field-scale detection to cross-domain deployment diagnostics or satellite-scale spatial prediction for any weed species, even where initial explorations of AI for herbicide reduction have demonstrated the potential of bridging these domains (León et al., 2024a). Two critical gaps exemplify this fragmentation. First, no deep learning detection model for *A. artemisiifolia* existed in the scientific literature prior to the present study, despite the species’ significance as one of the most aggressive and allergenic invasive weeds in temperate agriculture (Hasan et al., 2021). The only reported deep learning application to this species targets seed quarantine screening under controlled laboratory conditions (Liu et al., 2025), confirming the absence of any field-level detection capability. Without automated field detection, ragweed management relies on manual scouting—too slow and too expensive for the landscape-scale surveillance needed to intercept reproductive-stage plants before peak pollen dispersal, creating a critical bottleneck for source-reduction programs that aim to break the agricultural-to-urban dispersal chain documented above. Second, the individual AI components—detectors, embedding-based domain diagnostics, geostatistical interpolation, satellite-derived spectral predictors—have not been integrated into a coherent multi-scale workflow for any weed species, despite five decades of incremental progress in automated weed detection (Coleman et al., 2022). Even where detection models exist for other weed species, cross-domain transfer remains unreliable: models trained at one site frequently collapse when deployed in new geographies or cropping systems (Gao et al., 2024; Ilyas et al., 2023), and no standardized diagnostic exists to predict such failures before deployment. Meanwhile, citizen science platforms provide valuable occurrence records and species distribution models estimate invasion probability, but neither yields the quantitative per-unit-area density estimates that precision management requires.

This chapter addresses both gaps by presenting, evaluating, and releasing an open toolkit of AI tools for ragweed recognition and mapping at three complementary spatial scales. At the field scale, a YOLOv11 detection model is trained through a systematic hyperparameter sweep and deployed via sliced inference for high-resolution imagery (Section 3.1). At the cross-domain scale, ResNet50 embedding analysis quantifies the domain distance between training and deployment environments, and a two-phase experiment tests whether multi-domain training can recover the catastrophic failure predicted by those distances (Section 3.2). At the satellite scale, PRESTO foundation model embeddings are linked to drone-derived weed density through spatial statistical methods that account for geographic non-stationarity (Section 3.3). The emphasis throughout is on rigorous, transparent evaluation: each tool is tested against its failure modes, and performance boundaries are reported alongside achievements. Climate variability provides the unifying thread, because tools designed for non-stationary biological systems must themselves be adaptive—able to generalize across seasons, sites, and years as weed communities respond to changing conditions. The package source code and analysis workflows are released as a companion repository; the operational scripts and trained model weights are available from the corresponding author upon reasonable request.

The remainder of this chapter is organized as follows. Section 2 introduces the target species and the datasets assembled for training, embedding analysis, and satellite integration. Section 3 describes the methods at each spatial scale. Section 4 presents the results, following the same three-scale sequence. Section 5 synthesizes the findings, discusses their implications for weed management under climate variability, and acknowledges limitations. Section 6 distills five principal conclusions and identifies directions for future work.

## 2 Study System

### 2.1 Target Species: Ambrosia artemisiifolia

Common ragweed (*Ambrosia artemisiifolia* L.; Asteraceae) is an annual broadleaf weed native to North America that has become one of the most ecologically and economically significant invasive plant species worldwide. The species reproduces exclusively by seed, germinating in spring and completing its life cycle by late autumn. At the seedling stage—the phenological window most relevant to early-season detection— ragweed produces opposite pairs of deeply lobed, pinnately divided leaves that darken from light green to grey-green as the plant matures; leaves become alternate on the upper stem as the plant grows. These finely dissected leaves are the primary morphological cue for visual identification, yet they present a substantial classification challenge when observed at low spatial resolution: from a distance of two meters or greater, the compound leaf architecture collapses into an undifferentiated green cluster that is difficult to distinguish from co-occurring broadleaf species.

Four plant species were detected in the seedling-phase study system. In addition to ragweed (hereafter AMBEL, the standard EPPO identification code for *A. artemisiifolia*), the detection framework included the host crop lentil (*Lens culinaris* Medik.; LENCU), whose compound leaves contrast with ragweed at close range but converge to similar green patches at drone altitude; and two co-occurring weed species: prostrate knotweed (*Polygonum aviculare* L.; POLAV), a low-growing mat-forming species with small elliptic leaves; and pale persicaria (*Polygonum persicaria* L.; POLPE), an erect annual with lanceolate leaves. The morphological convergence between AMBEL and LENCU at drone altitude proved to be a central finding of this study and is discussed in detail in Section 4.1.

Ragweed’s significance extends well beyond crop yield reduction. In soybean, *A. artemisiifolia* interference can reduce yields by up to 83.7% while simultaneously impairing nitrogen fixation (Hall et al., 2021). The species is also among the most potent sources of allergenic pollen in temperate regions—a threat now monitored by deep learning–based pollen forecasting systems (Čorić et al., 2023)—and its range is expanding poleward under contemporary climate warming. Experimental evidence indicates that elevated atmospheric CO_2_ concentrations amplify ragweed biomass by up to 189% in urban environments, where the CO_2_ island effect is strongest (Ziska et al., 2003). Projections suggest that ragweed pollen sensitization in Europe will more than double by 2050 under moderate climate scenarios (Lake et al., 2017), with associated healthcare costs estimated in the billions of euros annually (Schaffner et al., 2020). In South America, ragweed pollen has been documented in aerobiological monitoring stations across multiple cities (Cherrez-Ojeda et al., 2024), and at least seven *Ambrosia* species are now recognized in the northwest of the continent, with *A. artemisiifolia* confirmed as introduced in Chile (Espinoza-Maticurena et al., 2025). Ecological niche modelling reveals that ragweed’s niche expansion in South America (expansion index = 0.407) exceeds that observed on any other continent (Song et al., 2023). Critically, agricultural populations of ragweed serve as source populations that feed urban pollen loads through wind dispersal—approximately 90% of pollen grains are deposited within 100 m of the source (Katz & Batterman, 2019)—creating a dual management imperative that links in-field weed control to public health outcomes (Hammad et al., 2026) and underscoring the need for field-level detection.

Despite the species’ agricultural and public health significance, no deep learning detection model for *A. artemisiifolia* existed prior to this work. Although hyperspectral imaging has been shown to discriminate ragweed from morphologically similar *Artemisia vulgaris* under controlled conditions (Dammer et al., 2013), this approach requires specialized sensors and has not been scaled to field-level deployment. More broadly, the gap reflects several interacting factors: the morphological similarity of broadleaf weed seedlings makes species-level annotation laborious; labeled training datasets for ragweed are scarce relative to agronomic staples; and the species has not been prioritized in the precision agriculture literature, which has focused predominantly on cereal and row-crop weed communities. The absence of automated detection is notable given that ragweed has been deemed important enough to warrant dedicated citizen science surveillance platforms (Dirr et al., 2025) and that genomics-informed species distribution models reveal distinct invasion potentials among different source populations (Putra et al., 2024)—both findings that would benefit from field-level validation data. The present chapter addresses this gap directly by training, evaluating, and releasing the first dedicated ragweed detection model.

### 2.2 Study Sites and Datasets

The primary study site was the Santa Rosa lentil (*Lens culinaris*) paddock, a 3.42 ha commercial field located near Retiro in the Maule Region of central Chile (36.532*^◦^*S, 71.913*^◦^*W; EPSG:32719, UTM Zone 19S). The paddock was surveyed during the early growing season (September 10–17, 2024) using a DJI unmanned aerial vehicle (UAV) operated at approximately 2 m flight altitude, yielding a ground sampling distance of approximately 0.5 mm per pixel. A total of 1,685 photograms were acquired across the survey, within which four weed species were detected: AMBEL, LENCU, POLAV, and POLPE. The Santa Rosa site served as the anchor for all three analytical threads of this chapter: field-scale detection, geostatistical mapping, and satellite-scale integration.

Ground-level smartphone surveys were conducted at three additional sites in the same region to evaluate model generalization to field-collected imagery. Ambrosia Santa Rosa comprised 75 images yielding 1,390 detections; CATO Maiz comprised 67 images with 2,443 detections; and Trigo Corregidas comprised 48 images with 1,372 detections. Across the three sites, 190 images produced 5,205 detections, with 87% of images (165 of 190) containing valid GPS coordinates in their EXIF metadata.

Training data for the detection model (Table 1) were drawn from two Chilean annotation campaigns. The CL_Seba dataset comprised 2,065 Roboflow-hosted images with bounding-box annotations, augmented offline with a 3*×* strategy (random cropping 0–29%, rotation *±*15*^◦^*, and salt-and-pepper noise) to produce 4,955 training tiles. The CL_Alberto dataset contributed an additional 659 independently annotated images from a separate Santa Rosa transect. A third Chilean source, CL_StaRosa (75 images), remained unlabelled and was used exclusively in the embedding analysis.

**Table 1.**
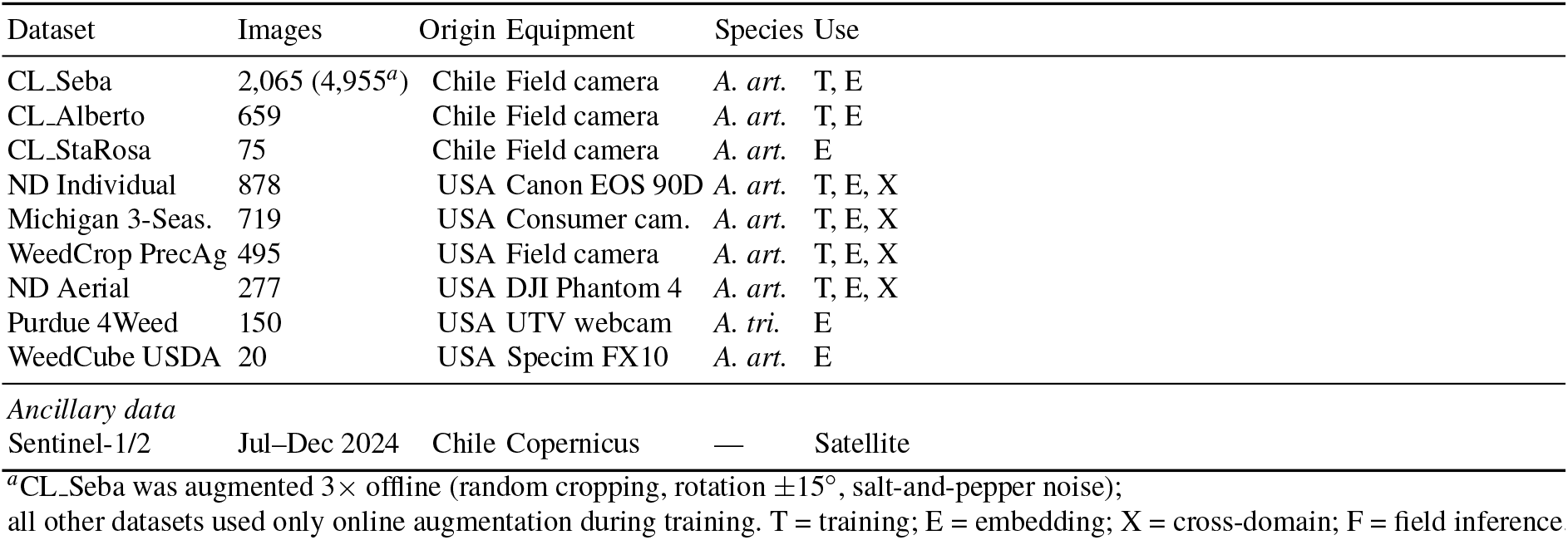
Summary of datasets used in training, embedding analysis, and cross-domain experiments. Origin indicates country or region of data collection. Use column specifies participation in training (T), embedding analysis (E), cross-domain benchmark (X), or field inference (F).

For the cross-domain generalization experiments, six international databases were assembled from publicly available repositories. ND Individual contained 878 images captured with a Canon EOS 90D in North Dakota, USA. Michigan 3-Season provided 719 images collected across three growing seasons (2021–2023) using consumer cameras. WeedCrop PrecAg contributed 495 images spanning eight crop-background categories from multiple US field sites. ND Aerial comprised 277 images acquired with a DJI Phantom 4 Pro in North Dakota. Purdue 4Weed contained 150 images captured by a webcam mounted on a utility terrain vehicle; this database targets *Ambrosia trifida* (giant ragweed), a morphologically distinct congener, and was therefore included in the embedding analysis but excluded from the detection benchmark. WeedCube USDA provided 20 pseudo-RGB images rendered from hyperspectral cubes (Specim FX10, 224 spectral bands, 400–1,000 nm), likewise included in embedding analysis only.

In total, the embedding space analysis encompassed 5,338 images from nine databases spanning two continents (2,799 Chilean and 2,539 international). The cross-domain training experiment drew on 7,981 images (CL_Seba: 4,955; CL_Alberto: 659; international training partition: 2,367). A summary of all datasets is provided in Table 1.

Ancillary data for the satellite-scale analysis included Sentinel-1 and Sentinel-2 imagery for the temporal window July–December 2024 was accessed via the Copernicus Data Space and used as input to the PRESTO foundation model (Section 3.3.1).

## 3 Methods

### 3.1 Field-Scale Detection

#### 3.1.1 Object Detection Architecture

Adult-phase detection training (December 2024 ground-level surveys). Single-class detection of common ragweed (*Ambrosia artemisiifolia*; AMBEL) was performed using YOLOv11l, the large variant of the YOLOv11 architecture (Jocher et al., 2024), which comprises 25.3 million parameters across 464 fused layers and operates at 86.6 GFLOPs. The backbone was initialized from weights pretrained on the COCO benchmark, providing general visual feature representations prior to domain-specific fine-tuning.

The training dataset was sourced from a Roboflow-hosted repository of 2,065 tiles annotated at 2048 *×* 2048 px native resolution, cropped to approximately 1050 *×* 915 px during Roboflow preprocessing, and licensed under CC BY 4.0. To identify the most effective training configuration, a four-run hyperparameter sweep was conducted (Table 2). All runs shared a common optimization backbone: stochastic gradient descent (SGD) with an initial learning rate of 0.01, linearly decayed to a final learning rate of 0.0001 (final ratio lrf = 0.01). Learning rate warmup was applied over the first 3 epochs, with momentum ramping from 0.8 to 0.937 and bias learning rate starting at 0.1. All runs used momentum of 0.937, weight decay of 0.0005, mosaic augmentation, and an early-stopping patience of 15 epochs. All runs used a fixed random seed of 42 with deterministic mode enabled, promoting highly reproducible training given identical hardware, software, and driver configurations.

**Table 2.** YOLOv11l training configuration across four experimental runs. All runs used SGD with lr_0_ = 0.01, linear decay to lr *_f_* = 0.0001, momentum 0.937, seed 42, patience 15, and deterministic mode.

| Parameter | Run 1 | Run 2 | Run 3 | Run 4 |
| --- | --- | --- | --- | --- |
| Dataset | Seba 640 | Seba Aug. | Seba Aug. | Combined |
| Images (train/val/test) | 1,450/411/204 | 4,335/411/209 | 4,335/411/209 | 4,911/466/237 |
| Transfer from | COCO | COCO | COCO | Run 3 |
| Image size (px) | 640 | 1024 | 640 | 640 |
| Batch size | 32 | 8 | 64 | 64 |
| GPU(s) | 1 × 4090 | 1 × 4090 | 2 × 4090 | 2 × 4090 |
| Epochs (completed) | 100 | 100 | 100 | 16 |
| Training time (min) | 26.2 | 164.0 | 39.9 | 7.5 |

Run 1 served as the baseline, training on 1,450 images at 640 px with a batch size of 32 on a single RTX 4090 GPU for 100 epochs (26.2 min). Run 2 introduced a 3*×* offline augmentation strategy applied to the same source imagery, expanding the training set to 4,335 images. Augmentations included random cropping (0–29%), rotation (*±*15*^◦^*), and salt-and-pepper noise. The image resolution was increased to 1024 px while batch size was reduced to 8 to maintain memory feasibility on a single RTX 4090 GPU; this run required 164.0 min to complete 100 epochs. Run 3 retained the augmented 4,335-image dataset but returned to 640 px tiles and scaled batch size to 64 using two RTX 4090 GPUs in DataParallel mode. This configuration completed 100 epochs in 39.9 min, yielding a validation mAP50 of 0.886—the highest across all runs. The comparison between Run 2 and Run 3 demonstrates that batch size scaling through multi-GPU training outperforms resolution scaling for this detection task. Run 4 used Run 3 weights as initialization and trained on a combined dataset of 5,614 images drawn from two annotation sources; early stopping halted training at epoch 16.

The detection loss follows the YOLOv11 default configuration (Jocher et al., 2024), combining Complete Intersection over Union (CIoU) for bounding-box regression (weight = 7.5), binary cross-entropy for classification (weight = 0.5), and distribution focal loss (DFL) for localization quality estimation (weight = 1.5). Training was conducted with automatic mixed precision (AMP) enabled. The total training objective is:

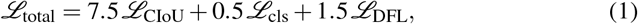

where *ℒ*_CIoU_ denotes the Complete Intersection over Union loss for bounding-box regression, *ℒ*_cls_ is the binary cross-entropy loss for single-class classification, and *ℒ*_DFL_ is the distribution focal loss for localization quality estimation.

Evaluation metrics follow the COCO protocol: mean average precision is computed at intersection-over-union thresholds of 0.50 (mAP_50_) and 0.50–0.95 in 0.05 increments (mAP_50:95_). For single-class detection, mAP_50:95_ is computed as:

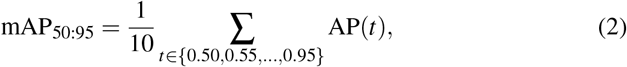

where AP(*t*) is the average precision at IoU threshold *t*. During validation, the confidence threshold for candidate predictions was set to 0.001 with non-maximum suppression (NMS) at IoU = 0.7.

A critical deployment parameter was the per-image maximum detection count (max_det), which was raised from the COCO default of 300 to 10,000. Dense weed infestations routinely produce hundreds to thousands of detections per image; the default cap would truncate results and artificially suppress recall in high-density fields. This parameter is frequently overlooked in YOLO implementations but is critical for accurate counting in dense agricultural populations.

#### 3.1.2 Slicing Aided Hyper Inference

Adult-phase field inference (December 2024). Ground-level smartphone images acquired during field surveys were captured at 4032 *×* 3024 px, substantially exceeding the 640 px tile resolution on which the detector was trained. Submitting full-resolution images directly to the model results in severe downscaling and the loss of fine spatial detail necessary to resolve individual AMBEL plants. To address this resolution mismatch, inference was performed using Slicing Aided Hyper Inference (SAHI) (Akyon et al., 2022), which partitions each image into overlapping sub-tiles at the native training resolution before aggregating predictions.

SAHI was configured with a slice size of 640 px to match the training resolution exactly, an overlap ratio of 0.20 to ensure continuity across slice boundaries, and a confidence threshold of 0.25. Non-maximum suppression (NMS) was applied across slice boundaries to merge duplicate predictions arising from the overlapping regions, using the library default IoU threshold of 0.50 for the postprocess matching step. The pipeline was deployed across 3 field sites, processing 190 images and yielding 5,205 total detections. Of the 190 images, 165 (87%) contained valid GPS coordinates embedded in EXIF metadata; geographic coordinates were extracted from these tags and used to produce a georeferenced point shapefile exported in EPSG:4326. Per-image detection counts ranged from 1 to 94, reflecting substantial spatial variation in AMBEL density across sampled locations.

#### 3.1.3 Drone-Scale Multi-Model Comparison

Seedling-phase drone evaluation (September 2024). To evaluate detection behavior at drone operational altitude (*≈*2 m, GSD *≈*0.5 mm/pixel), a stratified sample of ten drone images was selected from the 591 photograms acquired over the Santa Rosa paddock: three low-density scenes, four medium-density scenes, and three high-density scenes, determined by preliminary detection counts. Three independently trained model variants were compared: (i) the Original model (YOLOv11x, 56.9M parameters, trained at 1024 px, SAHI slice size 1024 px); (ii) the Intermediate model (YOLOv11l, 25.3M parameters, trained at 2048 px, SAHI slice size 2048 px); and (iii) the Retrained model (YOLOv11l, 25.3M parameters, trained at 2048 px with a different random seed, SAHI slice size 2048 px). All three models achieved comparable validation mAP_50_ on their respective training partitions (0.846–0.868). The evaluation focused on two metrics: total detection counts per image and per-species class distribution (AMBEL, LENCU, POLAV, POLPE), with particular attention to the combined broadleaf fraction (AMBEL + LENCU) as an indicator of species-level discrimination reliability at altitude. Confidence threshold was set to 0.25 with NMS IoU of 0.50 for all three models.

#### 3.1.4 Geostatistical Density Mapping

Kriging analysis was performed on seedling-phase drone detections (September 2024). Point-level detection counts were interpolated to continuous density surfaces using ordinary kriging. The interpolation domain was defined by the Santa Rosa lentil paddock (Section 2.2), discretized to a regular 5 m grid of 51 *×* 32 cells. In addition to AMBEL, three co-occurring species were mapped under the same framework: the host crop lentil (*Lens culinaris*; LENCU), prostrate knotweed (*Polygonum aviculare*; POLAV), and spotted lady’s-thumb (*Polygonum persicaria*; POLPE). Estimated density ranges across the paddock were 1.9–393.4 plants per cell for AMBEL, 27.4–402.4 for LENCU, 0.4–5.3 for POLAV, and 0.1–60.3 for POLPE.

Ordinary kriging was selected over universal kriging because the interpolation domain covers a single paddock of 3.42 ha where large-scale polynomial trends are unlikely to dominate; under these conditions the constant-mean assumption provides a more parsimonious model and avoids overfitting a trend surface. Spatial dependence was modeled using an exponential semivariogram fitted by least squares. Interpolation was performed in SmartMap geostatistical software (CENIA, Chile).

The spatial structure of detection counts was characterized via the exponential semi-variogram model. For lag distance *h >* 0, the semivariance is:

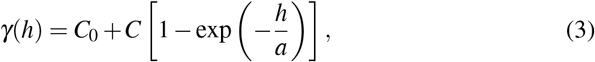

where *C*_0_ is the nugget variance (measurement error and micro-scale variation), *C* is the partial sill (structured variance), and *a* is the practical range. For the AMBEL layer, the fitted model yielded total sill *C*_0_ + *C* = 86,133, range *a* = 81.2 m, and nugget-to-sill ratio *C*_0_*/*(*C*_0_ + *C*) = 0.50, indicating that half the variance arose from unresolved small-scale heterogeneity and half from structured spatial correlation.

The resulting continuous density surfaces were subsequently used as ground-truth reference layers for the satellite-scale analysis presented in Section 3.3.

### 3.2 Cross-Domain Generalization

#### 3.2.1 Embedding Space Analysis

Adult-phase cross-domain analysis. Prior to any cross-domain training, the feature-space relationship between the Chilean and international databases was characterized using a frozen convolutional encoder, so that observed domain gaps could be quantified independently of any detection-specific fine-tuning.

##### Feature extraction

All images were passed through a ResNet-50 backbone (He et al., 2016) loaded with ImageNet V2 pretrained weights. The final classification layer was discarded; embeddings were read from the global average pooling layer, yielding one 2,048-dimensional vector per image. Each image was resized to 224 *×* 224 pixels and normalized with the ImageNet channel means and standard deviations. No data augmentation was applied during this step; the deliberate omission preserves the acquisition-specific characteristics (sensor, altitude, illumination) that distinguish each database, so that the measured distances reflect real deployment conditions rather than artificially equalized inputs.

##### Dataset coverage

Embeddings were computed for 5,338 images drawn from nine databases spanning two continents: 2,799 Chilean images and 2,539 international images. The three Chilean databases were CL_Seba (2,065 images, Roboflow-augmented field captures), CL_Alberto (659 images), and CL_StaRosa (75 unlabelled images from Santa Rosa). The six international databases were ND Individual (878 images, Canon EOS 90D), Michigan 3-Season (719 images, three growing seasons 2021–2023), WeedCrop PrecAg (495 images, eight crop-background categories), ND Aerial (277 images, DJI Phantom 4 Pro), Purdue 4Weed (150 images, webcam mounted on a utility terrain vehicle; note that this database targets *Ambrosia trifida*, giant ragweed, rather than *A. artemisiifolia*), and WeedCube USDA (20 hyperspectral pseudo-RGB images). Multi-class databases were filtered through a ragweed whitelist before embedding, retaining only images that contained at least one *Ambrosia* annotation.

##### Dimensionality reduction and domain-distance metrics

Embeddings were first reduced from 2,048 to 50 dimensions via Principal Component Analysis (PCA), and then projected to two dimensions with Uniform Manifold Approximation and Projection (UMAP) (McInnes et al., 2018) (*n_*neighbors = 15, min_dist = 0.1, cosine metric) and with t-SNE (Van der Maaten & Hinton, 2008) (perplexity = 30) for visual inspection.

Domain distance was formalized using the Maximum Mean Discrepancy (MMD) (Gretton et al., 2012). Let **X** = *{***x**_1_*,…,* **x***_m_}* denote the 50-dimensional PCA-reduced embeddings from the Chilean pool, and **Y** = *{***y**_1_*,…,* **y***_n_}* the embeddings from an international database. The empirical MMD^2^ statistic is:

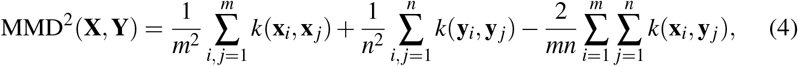

where 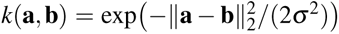 is the Gaussian radial basis function kernel and bandwidth *σ* is set to the median of all pairwise Euclidean distances (the median heuristic). Statistical significance was assessed by a permutation test with 1,000 resamples. Pairwise MMD values between the Chilean pool and each international database ranged from 0.27 to 0.43.

These values informed a four-tier deployment-gate framework that maps MMD ranges to recommended actions: direct deployment, augmentation and monitoring, active learning before deployment, and full retraining, in order of increasing domain distance. The specific thresholds delineating these tiers were calibrated empirically from the Phase 1 cross-domain experiment (Section 3.2.2) and are reported with the results in Section 4.2. Thresholds are dataset-dependent and should be recalibrated for new crop species or geographies. The observed MMD range of 0.27–0.43 placed all Chilean-to-international pairs in the intermediate tiers, providing a quantitative, pre-registered basis for the cross-domain training experiment described in Section 3.2.2.

#### 3.2.2 Two-Phase Cross-Domain Experiment

The embedding analysis generated a falsifiable hypothesis: the measured MMD range of 0.27–0.43 was predicted to produce measurable detection failure when the Chilean-only model is applied to international data, and a combined model trained on both domains was predicted to partially or fully recover that performance. Both predictions were pre-registered before any model was evaluated on the international test set.

##### Phase 1—evaluation of the Chilean model on international data

The best-performing Chilean model (Run 3, mAP_50_ = 0.886 on the Chilean test set; see Section 3.1.1) was applied without modification to a unified international test set comprising 2,367 images with 4,914 bounding-box annotations from four databases: ND Individual (876 images), Michigan 3-Season (719 images), WeedCrop PrecAg (495 images), and ND Aerial (277 images). Two databases present in the embedding analysis were excluded from the detection benchmark: Purdue 4Weed was omitted because it depicts *A. trifida*, a morphologically distinct congener whose inclusion would conflate inter-species variation with domain shift; WeedCube USDA was omitted because hyperspectral pseudo-RGB reconstructions lack bounding-box annotations in a format compatible with the detection pipeline.

Before evaluation, all four retained databases underwent format unification: class labels were remapped to a single ragweed class, Pascal VOC XML annotations were converted to YOLO format, and image filenames were prefixed with a database identifier to prevent collisions.

##### Phase 2—combined-domain training and evaluation

A combined training set of 7,981 images was assembled by joining the full Chilean training partition (CL_Seba: 4,955 images; CL_Alberto: 659 images) with the international training partition (2,367 images). The international data were split 80/10/10 into training, validation, and test subsets using stratified sampling per source database with a fixed random seed of 42, ensuring proportional representation of each international collection in every partition.

The combined model shared the same architecture as the Chilean-only baseline (YOLOv11l, 25.3 million parameters, COCO-pretrained weights) and was trained for 50 epochs with a batch size of 16 on a single NVIDIA RTX 4090, at an input resolution of 640 px (Table 3). All other hyperparameters were held constant relative to the Chilean-only run to isolate the effect of training-data composition.

**Table 3.** Cross-domain experiment configuration. Phase 1 evaluated the Chilean-only model on international data; Phase 2 trained a combined model on both domains.

| Parameter | CL-Only (Phase 1) | Combined (Phase 2) |
| --- | --- | --- |
| Architecture | YOLOv11l | YOLOv11l |
| Parameters | 25.3M | 25.3M |
| Training images | 4,955 (CL only) | 7,981 (CL + INTL) |
| Epochs | 100 | 50 |
| Batch size | 64 | 16 |
| Image size (px) | 640 | 640 |
| GPU(s) | 2× RTX 4090 | 1× RTX 4090 |
| Training time (min) | 39.9 | 52.5 |
| Pretrained | COCO | COCO |

The combined model was evaluated on two held-out sets: (a) the international test set defined in Phase 1, providing the primary cross-domain performance metric; and (b) the Chilean test set used in the original training experiment, serving as a regression check to detect catastrophic forgetting of source-domain knowledge.

##### Success criteria

Two quantitative criteria were specified prior to evaluation. The primary criterion required that the combined model achieve a higher mAP_50_ on the international test set than the Chilean-only model (Phase 1 baseline). The guard-rail criterion required that the combined model retain at least mAP_50_ *>* 0.836 on the Chilean test set, corresponding to a regression of no more than five percentage points relative to the Chilean-only model’s score of 0.886. This tolerance (approximately 6% relative degradation) reflects the operational requirement that deployment performance must remain within the precision envelope needed for density-based management decisions.

### 3.3 Satellite-Scale Integration

#### 3.3.1 PRESTO Foundation Model Embeddings

Seedling-phase satellite integration (September 2024 drone data, Santa Rosa). Foundation models pre-trained on satellite imagery—such as SatMAE (Cong et al., 2022), which learns spatial-spectral representations through masked autoencoding of multi-temporal Sentinel-2 scenes—have demonstrated strong transfer to downstream agricultural tasks. While location-aware embeddings such as SatCLIP (Klemmer et al., 2025) encode geographic coordinates directly, temporal approaches can capture crop phenological dynamics that static location encodings miss. This study adopted PRESTO (Pretrained Remote Sensing Transformer), a lightweight transformer architecture pre-trained on diverse remote sensing time series for general-purpose representation learning (Tseng et al., 2023). PRESTO was developed within the WorldCereal project to enable efficient crop monitoring across heterogeneous agro-ecological contexts without requiring task-specific fine-tuning. The model ingests multi-band, multi-temporal sequences and produces compact embeddings that encode seasonal spectral dynamics, making it suitable for characterizing vegetation states through time rather than relying on single-date snapshots.

The study site was the Santa Rosa paddock (Section 2.2), a commercial lentil field serving as the anchor for the satellite-scale integration analysis. Input data comprised Sentinel-1 and Sentinel-2 imagery acquired over the temporal window July 1– December 31, 2024 (six months), spanning the main phenological phases of the lentil growing season. Sentinel-2 Level-2A surface reflectance data were accessed via the Copernicus Data Space Ecosystem openEO API. Cloud and cloud-shadow masking was handled internally by the WorldCereal processing chain using the Sentinel-2 Scene Classification Layer (SCL), which excludes cloud shadow (class 3), medium- and high-probability cloud (classes 8–9), and thin cirrus (class 10); monthly composites were generated from the remaining clear observations. PRESTO was applied at the native Sentinel-2 spatial resolution of 10 m, yielding 128-dimensional embedding vectors per pixel. After applying the paddock boundary mask, 424 valid pixels fell within the field extent and were retained for processing.

To enable spatial alignment with the 5 m kriging grid derived from drone-based weed detections (Section 3.1.4), the 128-dimensional embeddings were spatially registered to that grid. Following alignment, 380 pixels with complete kriging coverage within the paddock boundary were used for all downstream statistical analyses. This alignment step was necessary because the kriged weed density surfaces were generated at 5 m resolution, whereas PRESTO embeddings were produced at 10 m; nearest-neighbor assignment was applied to propagate embedding values to the finer grid. Nearest-neighbor resampling was preferred over bilinear or cubic interpolation because weed detection counts are inherently discrete, and interpolating between count values would introduce ecologically meaningless fractional densities.

A key conceptual motivation for using temporal embeddings in this context is the spectral detectability of the target weed species. Above-ground broadleaf species such as *Ambrosia artemisiifolia* and the host crop *Lens culinaris* contribute directly to the mixed reflectance signal captured by satellite sensors, as their canopy biomass, leaf pigmentation, and growth phenology modulate the reflectance in visible and near-infrared wavelengths. This contrasts with root-parasitic species such as *Orobanche* spp., for which the satellite signal is only indirect, mediated by crop stress responses rather than the parasite’s own foliar expression. Consequently, temporal foundation model embeddings were hypothesized to capture phenological divergence between clean and weed-infested zones over the growing season.

#### 3.3.2 Dimensionality Reduction and Correlation

Prior to spatial analysis, the 128-dimensional PRESTO embedding space was compressed using Principal Component Analysis (PCA). PCA was computed on the 380 aligned pixels within the Santa Rosa paddock, with components ranked by descending explained variance. The first three principal components (PC1, PC2, PC3) were retained for analysis, as they captured the majority of embedding variance (see Section 4.3 for specific percentages). The 128-dimensional PRESTO representation proved to be strongly dominated by a low-dimensional structure.

The retained principal components were used as candidate predictors of weed infestation at the pixel scale. Pearson correlation coefficients were computed between each of the first ten PCs and the kriged weed density surfaces for each of the four weed species detected in the field. Kriged densities represented spatially continuous surfaces derived from 1,685 drone-based detection points processed using YOLOv11 (Section 3.1.2), subsequently interpolated to 5 m grids via ordinary kriging as described in Section 3.1.4. This correlation analysis served as an initial screening step to identify which embedding dimensions carried the strongest linear associations with ground-truth weed abundance before applying more computationally intensive spatial methods. Because both embedding components and kriged densities exhibit spatial autocorrelation (confirmed by the Moran’s I tests in Section 3.3.3), the Pearson coefficients reported here should be interpreted as descriptive summaries rather than independent-sample inferential tests; formal significance is assessed through the spatial methods in the following subsection.

#### 3.3.3 Spatial Statistical Methods

Four complementary spatial statistical methods were applied to characterize the relationship between satellite-derived embeddings and drone-measured weed densities. Together, these methods tested for spatial structure in the embedding-weed association, identified locally coherent clusters, quantified the spatial scale of co-variation, and assessed spatial non-stationarity in regression relationships. All spatial analyses were implemented in Python using the PySAL ecosystem: esda for global and local Moran’s I, libpysal for spatial weight construction, and mgwr for geographically weighted regression (Fotheringham et al., 2002); cross-variograms were computed with SciPy 1.15.

##### Bivariate Moran’s I (Anselin, 1995)

Global bivariate spatial autocorrelation was computed using bivariate Moran’s I to determine whether the spatial correlation between PRESTO embeddings and kriged weed density was structured—that is, whether neighboring pixels exhibiting high embedding values also tended to co-occur with high weed densities, beyond what would be expected under spatial randomness. A distance-band spatial weight matrix with a threshold of 15 m was constructed and row-standardized, reflecting the 5 m grid spacing and targeting the immediate spatial neighborhood. Statistical significance was assessed via 999 Monte Carlo permutations, with significance declared at *α* = 0.05. Bivariate Moran’s I measures global spatial correlation between two variables across neighboring spatial units:

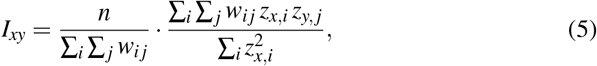

where *n* = 380 is the number of pixels within the paddock boundary, *w_i j_* is the row-standardized spatial weight (unity if the distance between pixels *i* and *j* is less than 15 m, zero otherwise), *z_x,i_* is the standardized PRESTO PC score at pixel *i*, and *z_y,j_* is the standardized kriged weed density at pixel *j*.

##### LISA cluster maps

Local Indicators of Spatial Association (LISA) were computed as the local decomposition of the global Moran’s I statistic, enabling cluster identification at the individual pixel level (Anselin, 1995). Each pixel was classified into one of four quadrant types: High-High (HH, high PC1 value co-located with high weed density), Low-Low (LL, low PC1 with low weed density), High-Low (HL, high PC1 with low weed density), or Low-High (LH, low PC1 with high weed density). Non-significant pixels were retained in the map but excluded from interpretation. The resulting cluster maps carry direct agronomic interpretation: HH pixels represent priority treatment zones where the satellite signal confirms elevated weed pressure; LL pixels indicate clean zones with concordant low spectral and weed signals; HL pixels are spectral outliers where embedding values are elevated but weed density is low; and LH pixels represent potentially missed infestations where weed density is high but the satellite signal does not flag it. The local form of Moran’s I at pixel *i* is:

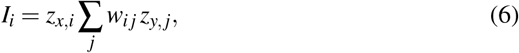

where *z_x,i_* is the standardized PRESTO PC1 at pixel *i* and *z_y, j_* is the standardized kriged weed density at neighboring pixel *j*. The sign of *I_i_* and the local weighted sum determine quadrant classification: HH when both are positive, LL when both are negative, HL when *z_x,i_ >* 0 but the neighbor sum is negative, and LH in the converse case.

##### Cross-variograms

Spatial cross-variograms were computed between PC1 and kriged weed density for each species to characterize the spatial scale at which satellite-weed co-variation reaches its maximum extent. Cross-variogram models were fitted to the empirical semi-variogram clouds, and the range parameter—the lag distance at which the semivariance stabilizes at the sill—was extracted as an estimate of the characteristic spatial scale of co-variation.

##### Geographically weighted regression

Geographically Weighted Regression (GWR) was applied to test for spatial non-stationarity in the relationship between satellite embeddings and weed density (Fotheringham et al., 2002). The dependent variable was the kriged weed density surface; predictors were PC1, PC2, and PC3. GWR allows regression coefficients to vary continuously across space, fitting a separate weighted regression at each pixel location using spatially decaying kernel weights. An adaptive bandwidth of 49 nearest neighbors was selected to account for the irregular effective sample density within the paddock boundary. The spatially varying coefficient surface was compared against a global Ordinary Least Squares (OLS) baseline fitted with identical predictors. Improvement in local fit (as assessed by local *R*^2^ values and AICc) was used to evaluate whether the embedding-weed relationship is spatially stationary or exhibits geographic variation that a global model would obscure. The predicted weed density at pixel *i* is:

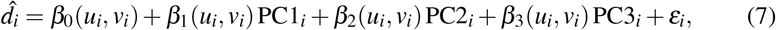

where (*u_i_, v_i_*) are the geographic coordinates of pixel *i*, *β_k_*(*u_i_, v_i_*) is the spatially varying coefficient for predictor *k*, and *ε_i_*is the local residual. Coefficients were estimated using an adaptive bisquare kernel with bandwidth equal to the 49th nearest neighbor, selected via AICc minimization.

#### 3.3.4 Single-Date Spectral Index Comparison

To benchmark the temporal foundation model approach against a simpler and operationally accessible alternative, a single-date spectral index analysis was performed using a Sentinel-2 image acquired on September 24, 2024—a date selected for its correspondence with peak phenological differentiation between weed-infested and clean zones during the lentil growing season. The image comprised 11 spectral bands (B2 through B12, excluding B1, B9, and B10), spanning the visible, red-edge, near-infrared, and shortwave infrared portions of the spectrum. All bands were resampled from their native Sentinel-2 resolutions of 10 m (visible and NIR: B2–B4, B8) and 20 m (red-edge and SWIR: B5–B7, B8A, B11–B12) to a common 5 m grid using nearest-neighbor interpolation, preserving original reflectance values without smoothing and matching the kriging grid resolution. After masking to the paddock boundary and restricting to pixels with valid kriging coverage, 1,503 pixels were available for analysis.

Nine spectral indices were computed from the resampled bands: the Normalized Difference Vegetation Index (NDVI), Enhanced Vegetation Index (EVI), Green Normalized Difference Vegetation Index (GNDVI), shortwave infrared difference index (SWIRd), Normalized Burn Ratio 2 (NBR2), Soil-Adjusted Vegetation Index (SAVI), Normalized Difference Moisture Index (NDMI), Bare Soil Index (BSI), and Normalized Difference Water Index (NDWI). These indices were selected to span vegetation greenness, canopy moisture, soil background effects, and canopy structure— dimensions relevant to distinguishing weed-affected from clean crop zones. Pearson correlation coefficients between each index and the kriged weed densities for all four species were then computed following the same procedure applied to the PRESTO principal components (Section 3.3.2).

The rationale for this comparison is that single-date NDVI and related indices represent the simplest operational pathway to satellite-assisted weed mapping, requiring no pre-training infrastructure, temporal data assembly, or embedding computation. If simple spectral indices derived from a single acquisition outperform or match temporal foundation model embeddings in terms of point-wise correlation with drone-measured weed density, this has substantive implications for recommending practical monitoring approaches in resource-limited agricultural contexts.

### 3.4 Toolkit Construction

The methods described in Sections 3.1–3.3 were implemented as a modular, installable Python package—the ragweed-ai-toolkit (v0.1.0-alpha, MIT license, Python *≥* 3.10)—released at https://github.com/agroia-lab/ragweed-ai-toolkit. The package is organized into eight submodules, each corresponding to a distinct analytical capability described in the preceding methods sections (Table 4).

**Table 4.** Toolkit module structure and correspondence to chapter methods sections. Each module implements the core algorithms described in the indicated section and exposes both a Python API and a command-line entry point.

| Module | Implements | Section | CLI entry point | Extras group |
| --- | --- | --- | --- | --- |
| detection | YOLOv11 training, SAHI, GPS extraction | 3.1.1, 3.1.2 | ragweed-train, -sahi | [detection] |
| embeddings | ResNet-50 features, MMD, UMAP/t-SNE | 3.2.1 | ragweed-embed, -mmd | [embeddings] |
| crossdomain | Multi-source datasets, transfer eval. | 3.2.2 | — | [detection] |
| orthomosaic | Drone mosaic tiling, active learning | 3.1.4 | ragweed-tile | [orthomosaic] |
| geostatistics | Semivariograms, ordinary kriging | 3.1.4 | ragweed-kriging | (core) |
| spatial | Moran’s I, LISA clusters, GWR | 3.3.3 | ragweed-lisa | [spatial] |
| satellite | PRESTO embeddings, spectral indices | 3.3.1, 3.3.4 | ragweed-indices | [satellite] |
| viz | Publication figures (300 DPI) | all | — | [viz] |

A key design objective was minimizing installation burden for users who require only a subset of the analytical capabilities. Heavy dependencies such as PyTorch, Ultralytics, and the Earth Engine API are isolated behind optional dependency groups (column “Extras group” in Table 4), so that, for example, a user interested only in spatial statistics can install the package with pip install -e “.[spatial]” without acquiring deep learning libraries. The core installation provides NumPy, pandas, GeoPandas, Matplotlib, SciPy, and scikit-learn—sufficient for geostatistical and visualization workflows.

Eight command-line entry points (Table 4) expose the principal workflows as standalone commands, enabling scripted batch processing without writing Python code. In addition, 27 CLI scripts organized into six pipeline-stage directories (01_detection through 06_satellite), available from the corresponding author upon request, provide thin wrappers around the package API with formatted terminal output. These scripts served as the operational interface during the analyses reported in Sections 4.1–4.3.

Reproducibility was addressed at multiple levels. All stochastic operations use fixed random seeds (default 42), and PyTorch deterministic mode is enabled during training. The test suite (7 modules, pytest) validates core functionality using synthetic data, ensuring that the package can be tested without access to proprietary imagery or GPU hardware. Six self-contained example scripts and four Jupyter notebook tutorials demonstrate complete workflows—from detection through spatial mapping—using synthetic data that ships with the repository. The notebooks are structured to mirror the three-act analytical sequence of this chapter: field detection, cross-domain analysis, and satellite integration.

The toolkit is released at alpha maturity (v0.1.0). The API may evolve as multi-site validation expands the range of tested conditions, but the analytical methods described in the preceding sections are stable and fully implemented.

## 4 Results

Table 5 summarizes the principal quantitative results across all three analytical scales before the detailed presentation that follows.

**Table 5.** Results at a glance: key metrics across the three analytical scales. Phase indicates seedling (S, September 2024) or adult (A, December 2024) phenological stage. Section references point to the detailed presentation.

| Scale | Key Metric | Value | Phase | Section |
| --- | --- | --- | --- | --- |
| Field | Best mAP<sub>50</sub> (AMBEL) | 0.886 | A | 4.1.1 |
| Cross-domain | CL model on INTL (collapse) | 0.108 | A | 4.2 |
| Cross-domain | Combined model on INTL (recovery) | 0.874 | A | 4.2 |
| Satellite | PRESTO PC1 × AMBEL ($r$) | 0.739 | S | 4.3 |
| Satellite | GWR $R^2$ (AMBEL / LENCU) | 0.882 / 0.910 | S | 4.3 |
| Satellite | Bivariate Moran's $I$ (PC1 × AMBEL) | 0.706 | S | 4.3 |
| Satellite | Single-date NDVI × AMBEL ($r$) | 0.837 | S | 4.3 |
| Satellite | Single-date NDVI × LENCU ($r$) | 0.890 | S | 4.3 |

### 4.1 Field-Scale Detection Performance

#### 4.1.1 Training Progression

Adult-phase training results (December 2024 data). The four-run training sweep revealed that data augmentation and batch size scaling were the dominant factors controlling detection performance, while input resolution and transfer learning from a domain-specific checkpoint yielded diminishing or negative returns (Table 6).

**Table 6.**
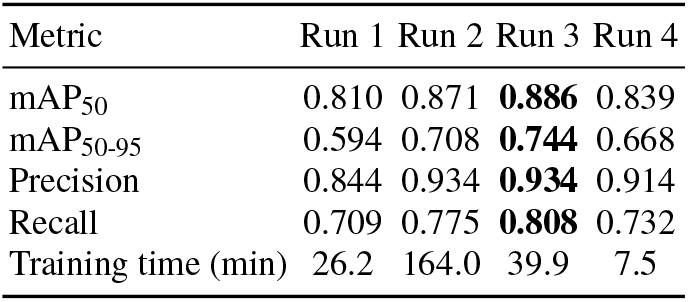
YOLOv11l training results across four experimental runs. Run 3 (bold) achieved the highest overall performance and was used as the production model. Run 4 used Run 3 weights as initialization (transfer learning).

| Metric | Run 1 | Run 2 | Run 3 | Run 4 |
| --- | --- | --- | --- | --- |
| mAP<sub>50</sub> | 0.810 | 0.871 | **0.886** | 0.839 |
| mAP<sub>50-95</sub> | 0.594 | 0.708 | **0.744** | 0.668 |
| Precision | 0.844 | 0.934 | **0.934** | 0.914 |
| Recall | 0.709 | 0.775 | **0.808** | 0.732 |
| Training time (min) | 26.2 | 164.0 | 39.9 | 7.5 |

Run 1 established a baseline of mAP_50_ = 0.810 with a precision of 0.844 and recall of 0.709, trained on 1,450 un-augmented images at 640 px with a batch size of 32 on a single RTX 4090 GPU (26.2 min). The limited recall indicated that the small, unaugmented training set was insufficient to capture the full range of ragweed appearance in field conditions.

Run 2 introduced the 3*×* offline augmentation strategy, expanding the training set from 1,450 to 4,335 images. This produced substantial gains: mAP_50_ increased to 0.871 (+7.5 percentage points over Run 1), and mAP_50-95_ rose from 0.594 to 0.708 (+19.2%). Precision reached 0.934. However, the image resolution was raised to 1024 px, which forced batch size down to 8 on a single GPU, extending training time to 164.0 min—a 6.3-fold increase relative to Run 1.

Run 3 retained the same augmented dataset but returned to 640 px and scaled batch size to 64 using two RTX 4090 GPUs in DataParallel mode. This configuration achieved the highest performance across all runs: mAP_50_ = 0.886, mAP_50-95_ = 0.744, precision of 0.934, and recall of 0.808. Compared with Run 2, Run 3 gained 1.7 percentage points in mAP_50_, 5.1 points in mAP_50-95_, and 4.3 points in recall, while completing training in 39.9 min—4.1 times faster. Run 3 achieved the highest mAP_50_ among all configurations (0.886), with training completed in 39.9 min (Table 6). The relationship between batch size and convergence is discussed in Section 5.1.

Run 4 attempted to extend Run 3 by fine-tuning its weights on a combined dataset of 5,614 images drawn from the CL_Seba and CL_Alberto annotation sources. However, the default learning rate of 0.01 proved too aggressive for transfer learning: the model peaked at epoch 1 (mAP_50_ = 0.839) and degraded steadily thereafter, triggering early stopping at epoch 16 (7.5 min total). This degradation pattern is interpreted in Section 5.1.

When all four models were applied via SAHI to nine held-out field images from three geographically distinct sites, Run 4 detected 58% more ragweed plants in the Santa Rosa images than Run 3 (122 versus 77 detections), despite its lower aggregate validation metrics. This gap between validation-set metrics and site-specific detection counts is discussed in Section 5.1.

#### 4.1.2 Drone-Scale Multi-Model Comparison

Seedling-phase drone comparison (September 2024). To assess detection behavior at drone operational altitude, three independently trained model variants were evaluated on a common set of ten drone images selected through stratified sampling by detection density (three low-density, four medium-density, and three high-density scenes). The three variants differed in architecture, training resolution, and SAHI slice size: the Original model (YOLOv11x, 56.9M parameters, trained and sliced at 1024 px), the Intermediate model (YOLOv11l, 25.3M parameters, trained and sliced at 2048 px), and the Retrained model (YOLOv11l, 25.3M parameters, trained and sliced at 2048 px with a different random seed). All three achieved comparable validation mAP_50_ on their respective training partitions (0.846–0.868), indicating that performance differences on the drone images were not attributable to model quality per se but rather to the interaction between resolution, slice configuration, and the morphological characteristics of the target species at altitude.

Table 7 summarizes the aggregate detection results. The Original model produced the most detections (6,306), attributable to its smaller SAHI slices generating more overlapping candidate tiles, while the two 2048 px models produced 4,359 and 4,653 detections respectively. However, the most consequential finding was not the total detection count but the systematic reversal of species labels between the resolution configurations.

**Table 7.** Drone-scale detection results across three model variants evaluated on ten stratified images. The class distribution reveals a systematic AMBEL/LENCU label inversion between the 1024 px and 2048 px configurations.

| Model (SAHI slice) | Total det. | AMBEL % | LENCU % | POLPE % |
| --- | --- | --- | --- | --- |
| Original (YOLOv11x @ 1024) | 6,306 | 11.6 | 65.4 | 17.8 |
| Intermediate (YOLOv11l @ 2048) | 4,359 | 52.2 | 31.5 | 15.0 |
| Retrained (YOLOv11l @ 2048) | 4,653 | 44.9 | 28.1 | 26.1 |

The Original model classified 65.4% of its detections as LENCU and only 11.6% as AMBEL. The 2048 px models inverted this ratio: the Intermediate model assigned 52.2% to AMBEL and 31.5% to LENCU; Retrained assigned 44.9% to AMBEL and 28.1% to LENCU. The same physical weeds in the same images received opposite species labels depending on the model configuration. This pattern held without exception across all ten images and all three density categories (Fig. 1).

**Fig. 1.**
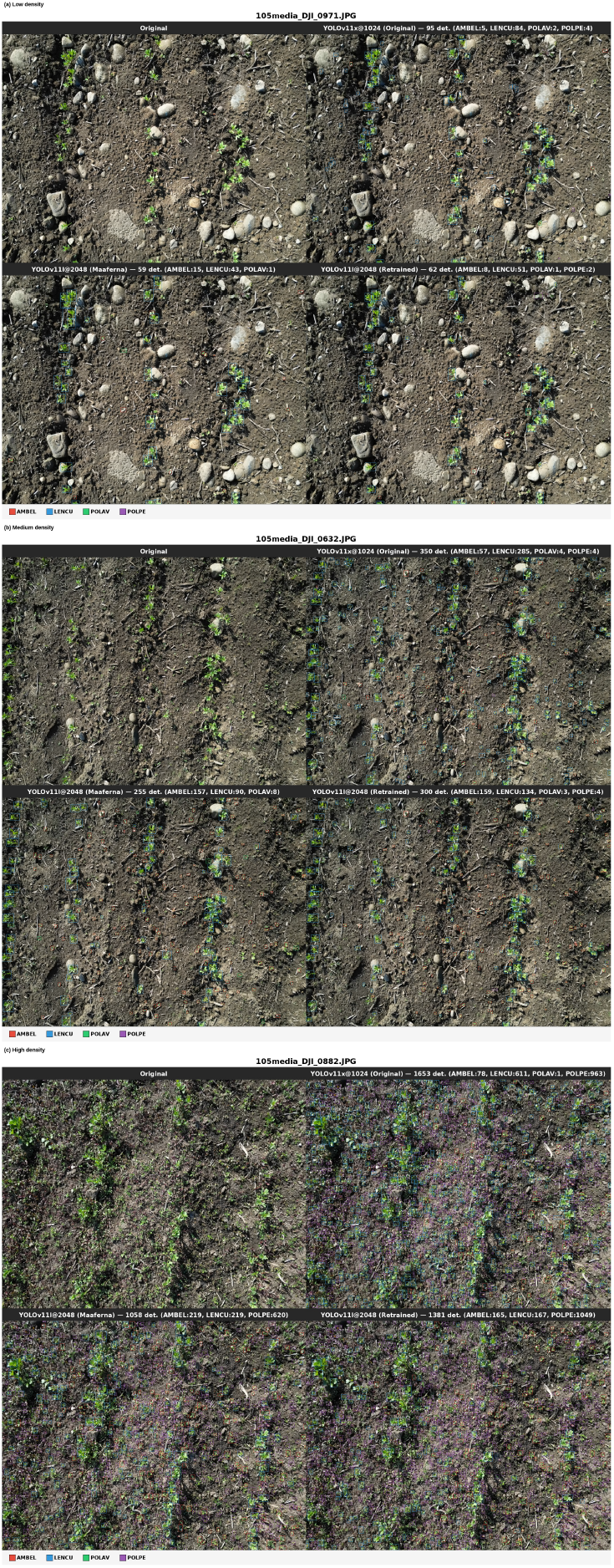
Multi-model weed detection comparison (seedling phase, September 2024) across three density categories in the Santa Rosa lentil paddock (*n* = 10 selected images from 591 drone photograms): (a) low density (DJI_0971, 62–95 detections), (b) medium density (DJI_0632, 255–350 detections), (c) high density (DJI 0882, 1,058–1,653 detections). Each scene shows four panels: original image (top left), YOLOv11x@1024 (top right), YOLOv11l@2048 Intermediate (bottom left), and YOLOv11l@2048 Retrained (bottom right). Bounding box colors: *A. artemisiifolia* (AMBEL, red), *L. culinaris* (LENCU, blue), *P. aviculare* (POLAV, green), *P. persicaria* (POLPE, magenta). SAHI sliced inference with confidence threshold 0.25 and NMS IoU 0.5. Flight altitude *≈*2 m (GSD *≈*0.5 mm/pixel). The AMBEL/LENCU label inversion between 1024 px and 2048 px configurations is visible across all density levels.

The combined AMBEL+LENCU fraction was considerably more stable across models than either species alone: 77.0% for Original, 83.7% for Intermediate, and 73.1% for Retrained. This consistency indicates that all three models reliably located broadleaf weed patches but disagreed on species identity within those patches. The mechanistic explanation for this inversion is developed in Section 5.5.

#### 4.1.3 Geostatistical Mapping

Point-level detection counts from the 1,685 drone photograms were interpolated to continuous density surfaces at 5 m resolution across the 3.42 ha Santa Rosa paddock using ordinary kriging, applied independently for each of the four weed species. Estimated density ranges were 1.9–393.4 detections per 5 m cell for AMBEL, 27.4–402.4 for LENCU, 0.4–5.3 for POLAV, and 0.1–60.3 for POLPE (Fig. 2). The kriged surfaces revealed pronounced spatial heterogeneity in weed pressure, with AMBEL and LENCU exhibiting complementary hot-spot distributions across the paddock. These continuous density surfaces served as the ground-truth reference layers for the satellite-scale analysis presented in Section 4.3.

**Fig. 2.**
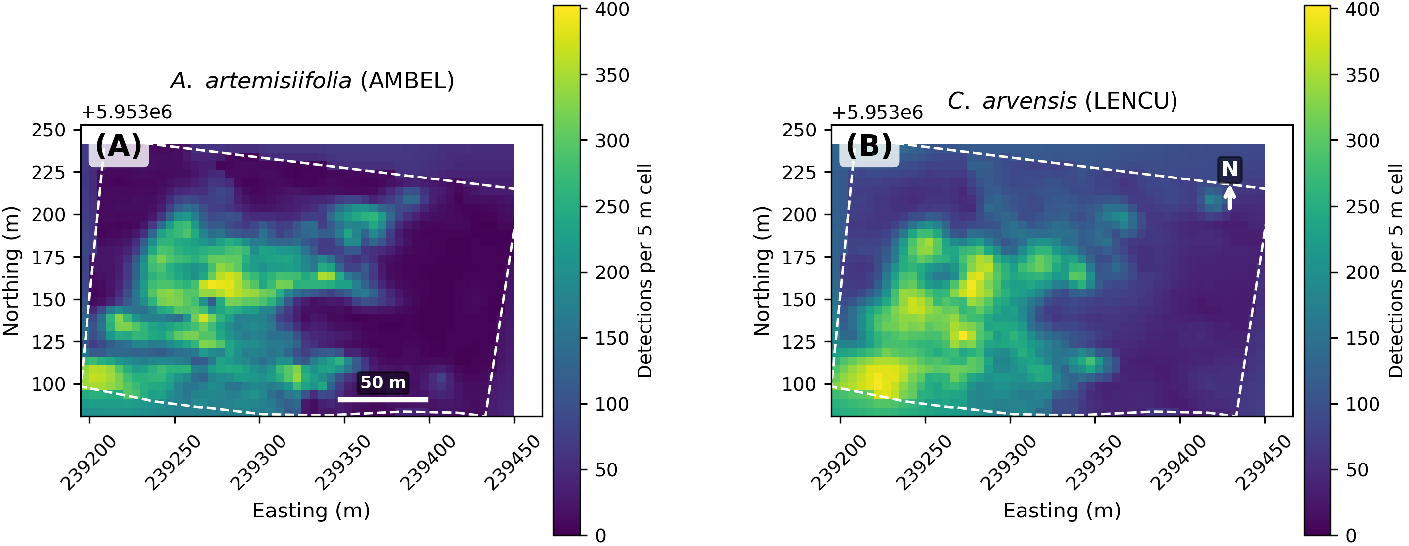
Kriged weed density surfaces (seedling phase) at 5 m resolution for (A) *A. artemisiifolia* (AMBEL, range 1.9–393.4 detections per cell) and (B) *L. culinaris* (LENCU, range 27.4–402.4 detections per cell) in the 3.42 ha Santa Rosa lentil paddock. Ordinary kriging with an exponential variogram model (sill = 86,133; range = 81.2 m; nugget/sill = 0.50) interpolated *n* = 1,685 georeferenced drone photograms onto a regular grid (51 *×* 32 cells). Viridis colormap; dashed white line indicates paddock boundary. Complementary hot-spot distributions are visible, with AMBEL concentrated in the eastern sector and LENCU more uniformly distributed.

### 4.2 Cross-Domain Generalization

#### Embedding space structure (adult phase)

The UMAP projection of 5,338 images from nine databases revealed that the embedding space is structured primarily by acquisition conditions rather than by species morphology (Fig. 3). Images clustered tightly by source database, with camera hardware, illumination regime, and background vegetation driving the dominant axes of variation. A clear geographic separation emerged between the 2,799 Chilean images and the 2,539 international images, confirming that field conditions in central Chile produce visually distinct imagery relative to North American sources.

**Fig. 3.**
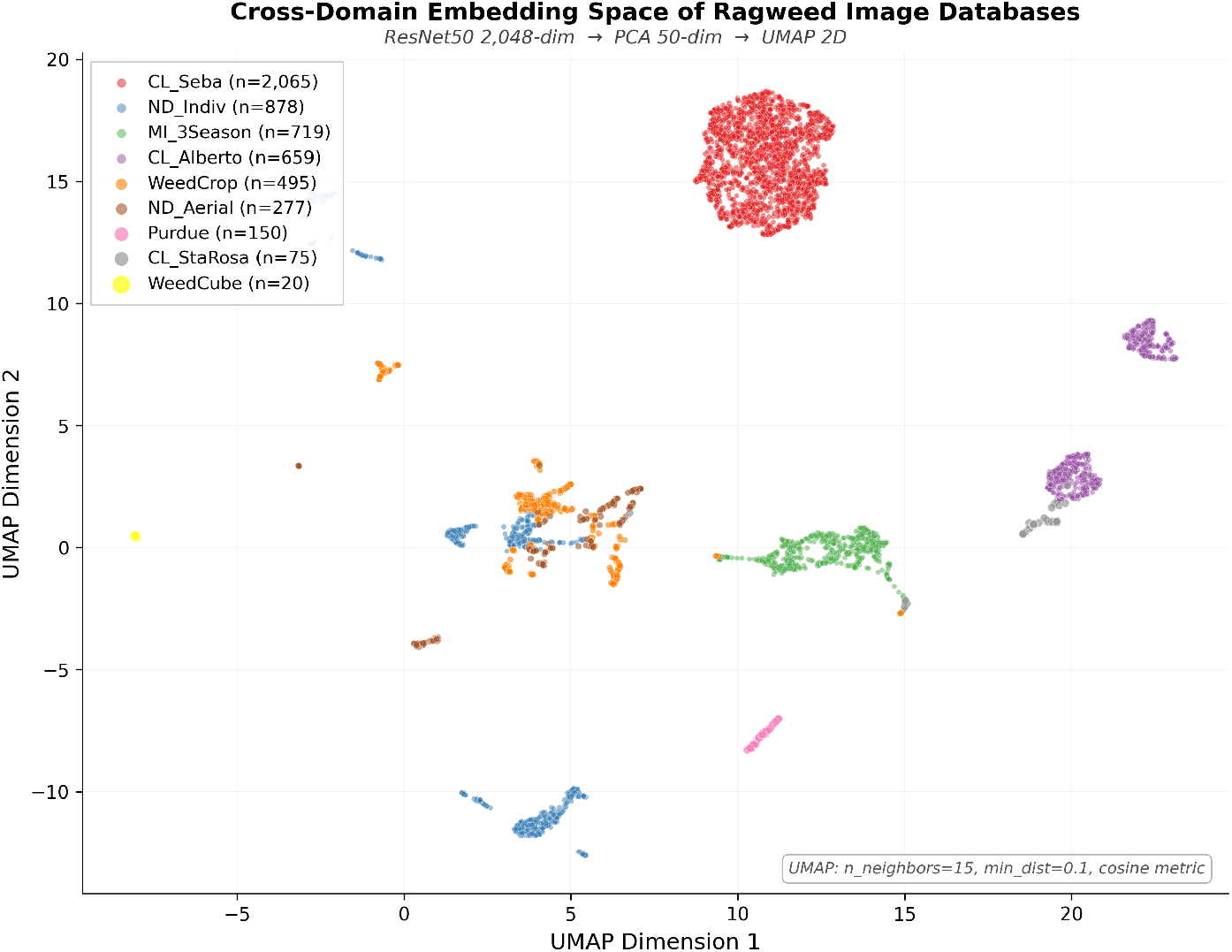
UMAP two-dimensional projection of ResNet50 embeddings (adult-phase cross-domain analysis) for *N* = 5,338 ragweed images across nine databases spanning two continents. Colors indicate database of origin: CL_Seba (*n* = 2,065), ND_Indiv (*n* = 878), MI 3Season (*n* = 719), CL_Alberto (*n* = 659), WeedCrop (*n* = 495), ND_Aerial (*n* = 277), Purdue (*n* = 150), CL_StaRosa (*n* = 75), Weed-Cube (*n* = 20). Embedding pipeline: ResNet50 2,048-dim *→* PCA pre-reduction to 50 dims *→* UMAP (*n*_neighbors_ = 15, min dist = 0.1, cosine metric, random state 42). Database-specific clustering dominates the feature space, confirming that domain (acquisition conditions) rather than species identity drives embedding separation.

Among the Chilean databases, CL_Seba (2,065 images) formed the densest single cluster, a consequence of Roboflow augmentation homogenizing the visual signature of the source field photographs. In the international group, ND Aerial (277 drone images) occupied an isolated region of the embedding space, reflecting the perspective shift between aerial and ground-level acquisition. WeedCube USDA (20 hyperspectral pseudo-RGB images) was the most distant outlier overall, consistent with its fundamentally different sensing modality. The databases exhibiting the greatest internal diversity—WeedCrop PrecAg (multiple crop backgrounds) and ND Individual (multiple growth stages)—were also the most internally dispersed in the projection, and their average MMD to all other databases was the lowest (0.272 and 0.267, respectively). In contrast, the most visually homogeneous databases exhibited the highest average MMD: WeedCube (0.518), Purdue (0.428), and CL_Seba (0.383). This pattern indicates that acquisition diversity within a database translates directly to proximity to other databases in the embedding space (Fig. 4).

**Fig. 4.**
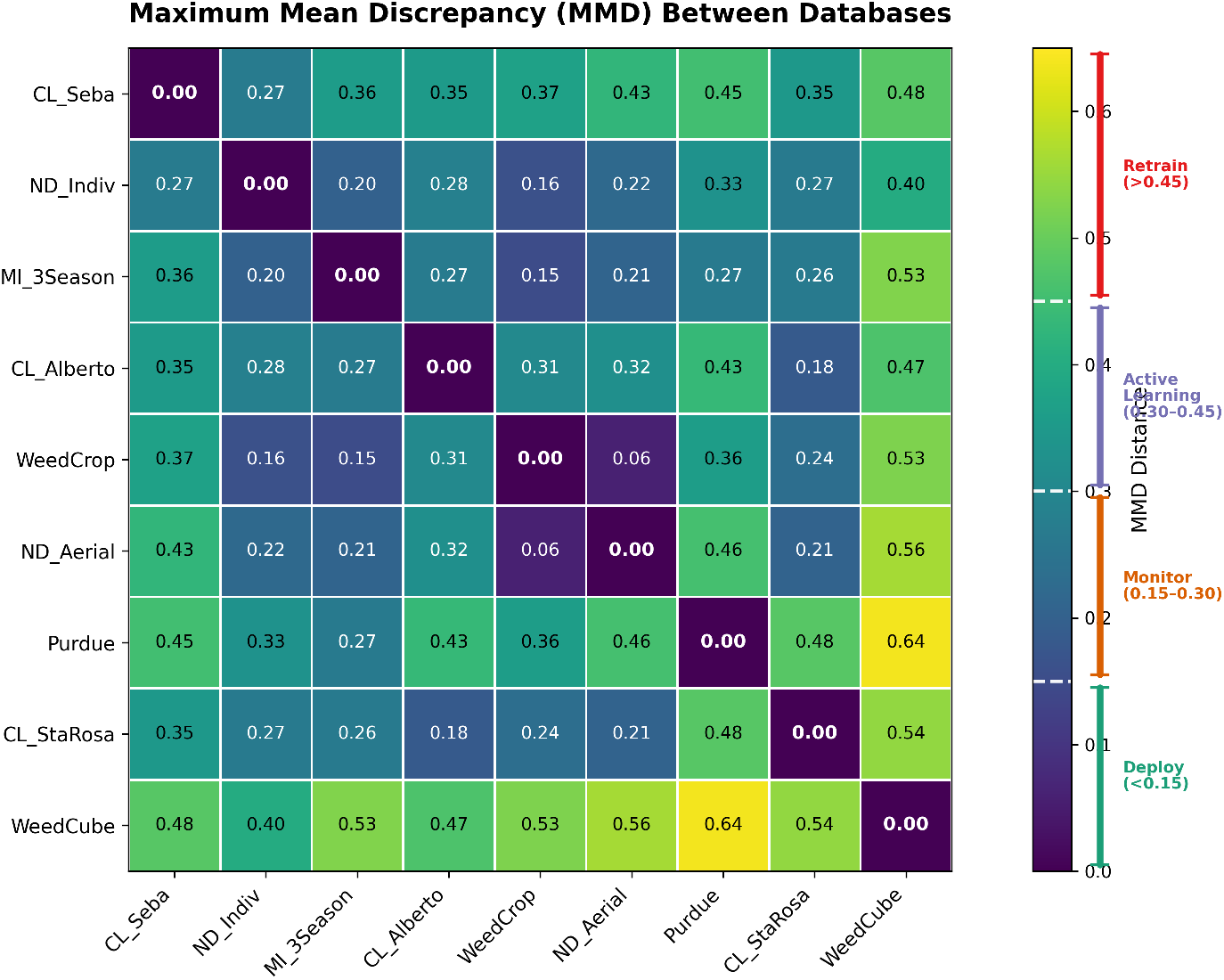
Pairwise Maximum Mean Discrepancy (MMD) distance matrix (adult-phase cross-domain analysis) across nine ragweed databases, computed with a Gaussian RBF kernel using median heuristic bandwidth on L2-normalized ResNet50 embeddings. Cell values report MMD distances (two decimal places). Chilean databases cluster together (MMD *<* 0.15); international curated databases are most distant (MMD = 0.27–0.43). WeedCube (*n* = 20) shows the highest distances (0.40–0.64), likely due to its small sample and controlled imaging conditions. Dashed lines on the colorbar indicate deployment gate thresholds: *<* 0.15 (deploy), 0.15–0.30 (monitor), 0.30–0.45 (active learning), *>* 0.45 (retrain). Database abbreviations: CL = Chilean, ND = North Dakota, MI = Michigan. Viridis colormap.

#### Phase 1: catastrophic cross-domain failure

The best-performing Chilean model (Run 3, mAP_50_= 0.886 on its Chilean test set) was applied without modification to the unified international test set of 2,367 images containing 4,914 bounding-box annotations from four databases. The results confirmed a near-complete detection failure: mAP_50_ dropped to 0.108, a decline of 77.8 percentage points (Table 8). Per-database analysis revealed that the model produced *zero* predictions across all four international databases at the evaluation confidence threshold of 0.001. This was not gradual degradation proportional to domain distance; it was a categorical inability to recognize ragweed in imagery from any international source. The severity of this failure is consistent with the MMD distances of 0.27–0.43 measured between the Chilean training pool and the international databases, which placed all pairs in the “active learning required” tier of the deployment-gate framework (Gretton et al., 2012).

**Table 8.** Cross-domain detection performance. Phase 1 evaluated the Chilean-only model on international data; Phase 2 evaluated the combined model on both domains. Precision and recall for the Chilean baseline are from the original training experiment (Section 3.1.1).

| Scenario | mAP<sub>50</sub> | mAP<sub>50-95</sub> | Precision | Recall |
| --- | --- | --- | --- | --- |
| CL model on CL test (baseline) | 0.886 | 0.744 | 0.934 | 0.808 |
| CL model on INTL test (Phase 1) | 0.108 | 0.047 | 0.268 | 0.180 |
| Combined on INTL test (Phase 2) | 0.874 | 0.635 | 0.835 | 0.772 |
| Combined on CL test (regression) | 0.742 | 0.435 | 0.757 | 0.660 |

#### Phase 2: multi-domain recovery

Training a combined model on 7,981 images drawn from both Chilean and international sources recovered international detection performance to mAP_50_ = 0.874, an improvement of 76.6 percentage points over the Phase 1 baseline (Table 8). This result satisfied the primary success criterion, which required that the combined model exceed the Chilean-only model’s international mAP_50_ of 0.108.

However, the combined model incurred a regression of 14.4 percentage points on the Chilean test set, with mAP_50_ declining from 0.886 to 0.742. This exceeded the pre-registered guard-rail tolerance of five percentage points (mAP_50_ *>* 0.836). The combined model differed from the baseline in several respects beyond training data composition: 50 epochs versus 100, batch size 16 versus 64, and a single GPU versus two—differences imposed by hardware memory constraints rather than experimental design. The training curve had not plateaued at epoch 50 (Fig. 5).

**Fig. 5.**
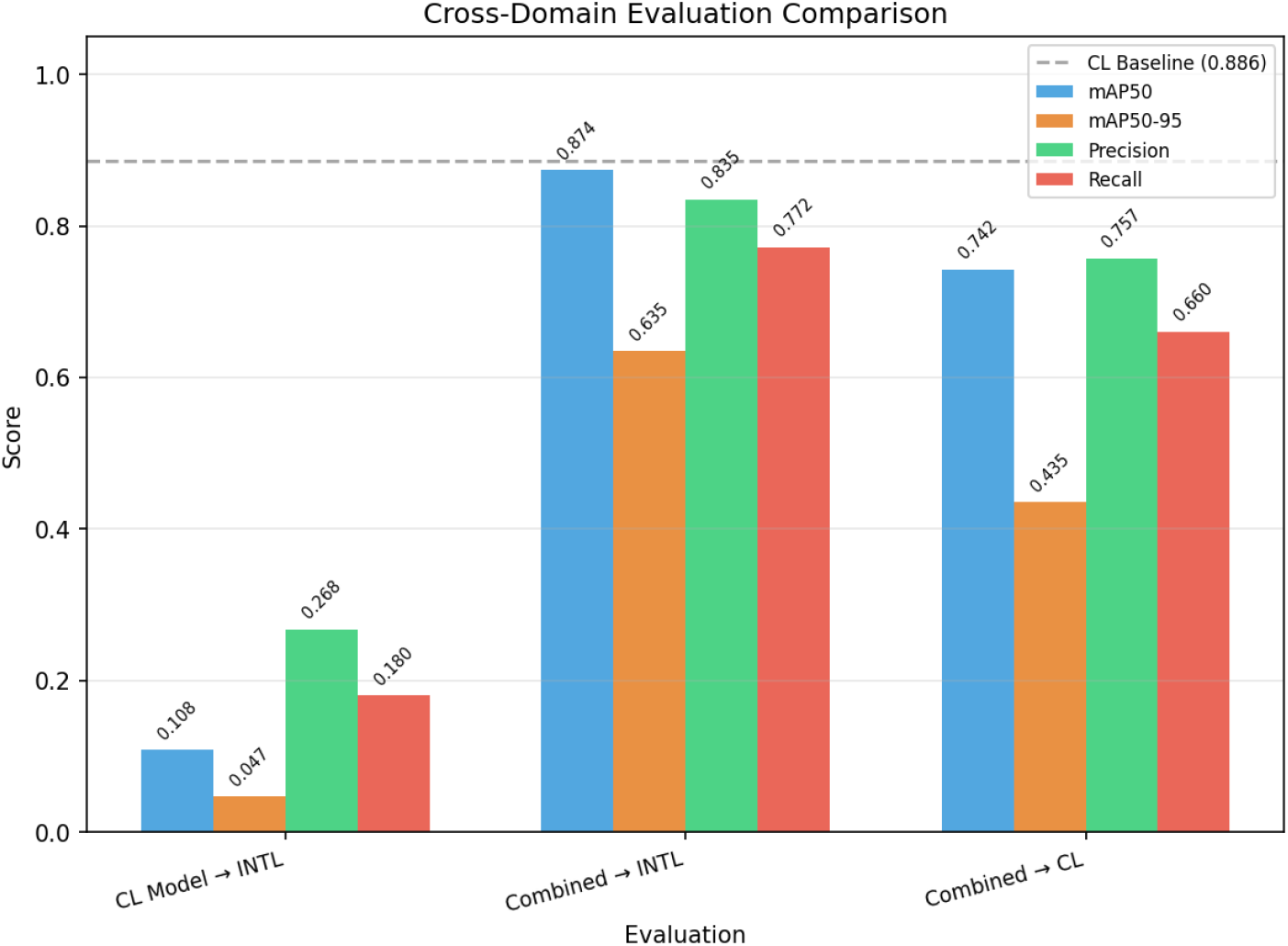
Cross-domain detection performance comparison (adult phase). Phase 1: Chilean-only model (Run 3, trained on *n* = 4,335 CL_Seba images, mAP_50_ = 0.886) evaluated on *n* = 2,367 international images from four databases (ND_Indiv, MI_3Season, WeedCrop, ND_Aerial), yielding mAP_50_ = 0.108 (effectively zero useful detections). Phase 2: combined model trained on *n* = 6,805 images (Chilean + international) recovered to mAP_50_ = 0.874 on international data (+76.6 percentage points) at the cost of moderate Chilean regression (0.886 *→* 0.742, *−*14.4 percentage points). mAP_50_ = mean Average Precision at IoU *≥* 0.50. Single-run evaluation; confidence intervals would require repeated training with different random seeds.

#### Five-model benchmark

To disentangle the contributions of dataset size, augmentation strategy, resolution, transfer learning, and domain diversity to cross-domain generalization, all five YOLOv11l models from the training progression were evaluated on a strictly held-out international test set of 235 images containing 512 ragweed annotations (Table 9). None of these images were used during training of any model.

**Table 9.** Five-model benchmark on 235 held-out international images (512 annotations). All models share the YOLOv11l architecture (25.3M parameters). Standard YOLO validation at 640 px, confidence threshold 0.001.

| Model | Training data | Imgs | mAP<sub>50</sub> | mAP<sub>50-95</sub> | P | R |
| --- | --- | --- | --- | --- | --- | --- |
| Seba 640 | CL_Seba | 2,065 | 0.173 | 0.080 | 0.367 | 0.221 |
| Lentejas 1024 | CL_Seba aug. | 4,955 | 0.056 | 0.023 | 0.185 | 0.127 |
| Lentejas 640 | CL_Seba aug. | 4,955 | 0.110 | 0.056 | 0.277 | 0.151 |
| CL Transfer | CL + Alberto | 5,614 | 0.169 | 0.080 | 0.316 | 0.240 |
| CL + Intl | CL + 4 intl | 7,981 | 0.791 | 0.507 | 0.696 | 0.752 |

The four Chilean-only models (Runs 1–4) achieved mAP_50_ values between 0.056 and 0.173, regardless of the strategy employed. Increasing the Chilean training set from 2,065 to 4,955 images through offline augmentation (Run 1 to Run 2) *decreased* performance from 0.173 to 0.056. Returning to 640 px resolution with larger batch size (Run 3) partially recovered to 0.110. Transfer learning from a second Chilean field (Run 4, 5,614 images) reached 0.169, still failing to exceed the smaller baseline. In contrast, Run 5—which added 2,367 international images to the Chilean pool—produced a phase transition: mAP_50_ jumped to 0.791, an increase of 62.2 percentage points over the best Chilean-only model. Domain diversity, not dataset size, was the distinguishing factor: the only model to achieve meaningful cross-domain detection included international training imagery.

#### Deployment gate validation

The embedding-based deployment-gate framework introduced in Section 3.2.1 specified four tiers of action based on pairwise MMD: values below 0.15 permit direct deployment, 0.15–0.30 warrant augmentation and monitoring, 0.30–0.45 require active learning before deployment, and values exceeding 0.45 necessitate full retraining (Gretton et al., 2012). The observed MMD range of 0.27–0.43 between Chilean and international databases placed all cross-domain pairs in the augmentation-required to active-learning-required tiers. Phase 1 confirmed this prediction: the Chilean-only model achieved near-zero detection on international imagery. Phase 2 validated the prescribed remedy: training with target-domain data recovered mAP_50_ to 0.874. The operational implications of this calibration are discussed in Section 5.4.

### 4.3 Satellite-Scale Integration

#### PCA structure and correlations (seedling phase)

Principal component analysis of the 128-dimensional PRESTO embeddings computed on 380 spatially aligned pixels revealed a strongly low-dimensional structure. The first three principal components captured 73.5% of the total embedding variance (PC1 = 42.7%, PC2 = 17.1%, PC3 = 13.7%), and ten components sufficed to explain 94.9%. PC1 emerged as the dominant linear predictor of weed density for the two principal broadleaf species: Pearson correlation coefficients were *r* = 0.739 (95% CI [0.690, 0.782]) for AMBEL and *r* = 0.717 (95% CI [0.664, 0.763]) for LENCU (*n* = 380; Fig. 6). PC1 alone accounted for 54.6% of AMBEL variance (*r*^2^ = 0.546) and 51.4% of LENCU variance (*r*^2^ = 0.514).

**Fig. 6.**
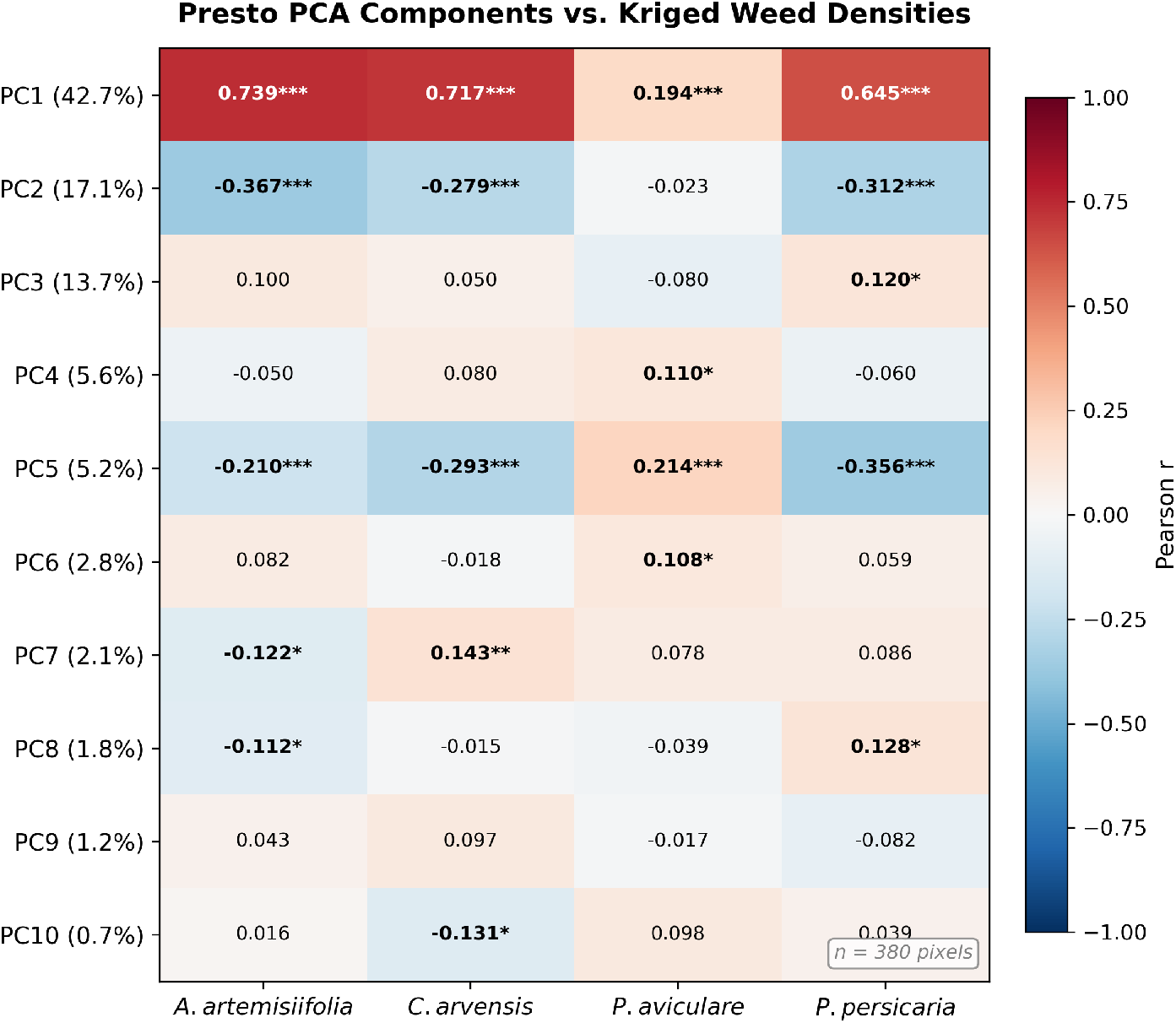
Pearson correlation heatmap (seedling phase) between PRESTO PCA components (PC1– PC10) and kriged densities of four species (*n* = 380 pixels, 999 permutations). PC1 (42.7% variance explained) shows the strongest associations with *A. artemisiifolia* (*r* = 0.739) and *L. culinaris* (*r* = 0.717). Higher-order components capture progressively weaker and species-specific signals. Asterisks denote *p <* 0.05 (*), *p <* 0.01 (**), and *p <* 0.001 (***).

Secondary components carried weaker but interpretable signals. PC2 showed moderate negative associations with AMBEL (*r* = *−*0.367) and POLPE (*r* = *−*0.312), while PC5 exhibited contrasting signs across species (POLAV: *r* = +0.214; LENCU: *r* = *−*0.293; POLPE: *r* = *−*0.356), suggesting that higher-order components encode subtler spectral contrasts that discriminate between species with distinct growth forms. POLAV (*P. aviculare*), however, showed no significant correlation with any spectral predictor (PC1: *r* = 0.194, 95% CI [0.095, 0.289], *n* = 380), reflecting its very low density range (0.4–5.3 plants per cell) and prostrate, low-biomass growth form, which provides insufficient canopy contribution for satellite-resolution detection (Fig. 7). This species-specific failure indicates that the spectral bridge does not extend to all weed functional types.

**Fig. 7.**
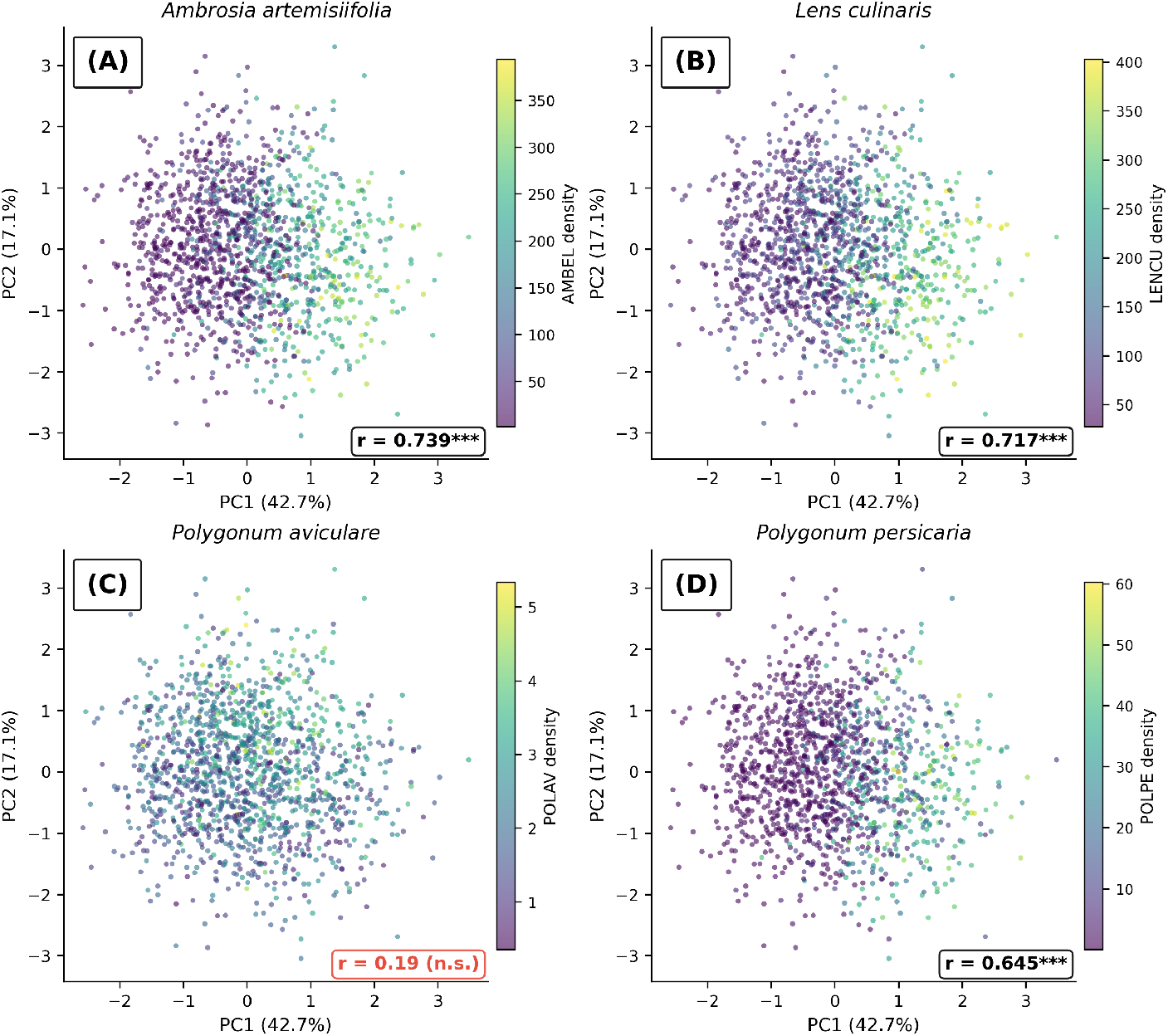
PCA scatter plots (seedling phase; PC1 vs. PC2) colored by kriging density for each species. (A) *A. artemisiifolia* (*r* = 0.739), (B) *L. culinaris* (*r* = 0.717), (C) *P. aviculare* (*r* = 0.194, n.s.), (D) *P. persicaria* (*r* = 0.645). PC1 and PC2 together explain 59.8% of Presto embedding variance. A clear density gradient along the PC1 axis is visible for *A. artemisiifolia* and *L. culinaris*. The absence of a density gradient for *P. aviculare* confirms that the spectral bridge is species-selective, limited to upright, biomass-dominant broadleaf species.

#### Single-date spectral index comparison

To contextualize the temporal embedding approach, Pearson correlations between kriged weed densities and nine spectral indices derived from a single Sentinel-2 acquisition (September 24, 2024) were compared against PRESTO PC1 and the full GWR model (Table 10).

**Table 10.** Comparison of satellite-based approaches for predicting weed density. Single-date indices were computed from Sentinel-2 imagery acquired on September 24, 2024 (*n* = 1,503 pixels at 5 m resolution). PRESTO embeddings integrate six months of multi-sensor observations (July–December 2024; *n* = 380 pixels at 10 m resolution). Best correlations per species are shown.

| Approach | AMBEL $r$ | LENCU $r$ | Notes |
| --- | --- | --- | --- |
| Single-date NDVI | 0.837 | 0.890 | Simple; highest point-wise $r$ |
| Single-date EVI | 0.771 | 0.829 | Reduced saturation bias |
| Single-date GNDVI | 0.787 | 0.853 | Green-band sensitive |
| PRESTO PC1 (128-dim, 6 months) | 0.739 | 0.717 | Temporal dynamics, multi-sensor |
| PRESTO GWR (PC1–PC3, local) | $R^2$: 0.882 / 0.910 | | Highest explanatory power |

Single-date NDVI achieved the highest point-wise correlations among all approaches, reaching *r* = 0.890 (95% CI [0.879, 0.900]) for LENCU and *r* = 0.837 (95% CI [0.821, 0.852]) for AMBEL (Table 10). EVI and GNDVI followed the same pattern at moderately lower magnitudes. PRESTO PC1, which integrates six months of multi-sensor observations into a single embedding axis, produced lower point-wise correlations (*r* = 0.739, 95% CI [0.690, 0.782] and *r* = 0.717, 95% CI [0.664, 0.763]), but when the spatial structure of the PRESTO embeddings was modeled through geographically weighted regression (see below), the local *R*^2^ reached 0.910 for LENCU and 0.882 for AMBEL. The mechanistic comparison between the two approaches is discussed in Section 5.2.

#### Bivariate spatial correlation

The preceding correlations treated each pixel as an independent observation. Bivariate Moran’s I was computed to test whether the embedding-weed association is spatially structured—that is, whether neighboring pixels with high embedding values systematically co-occur with high weed densities beyond what would be expected under spatial randomness (Anselin, 1995). A distance-band spatial weight matrix with a threshold of 15 m was constructed and row-standardized, and statistical significance was assessed via 999 Monte Carlo permutations.

The results confirmed strong bivariate spatial autocorrelation. PC1 *×* AMBEL yielded a Moran’s *I* = 0.706 (*p* = 0.001), and PC1 *×* LENCU yielded *I* = 0.694 (*p* = 0.001). These values are notably high for a bivariate statistic at 10 m resolution and indicate that the satellite-derived embedding gradient and the ground-truth weed density gradient are not merely correlated in aggregate but are spatially coherent across the paddock. Secondary components showed weaker, negatively signed spatial associations (PC2 *×* AMBEL: *I* = *−*0.321; PC2 *×* LENCU: *I* = *−*0.254; both *p* = 0.001), consistent with the opposing sign of the PC2 Pearson correlations reported above.

#### LISA cluster analysis

Local Indicators of Spatial Association (LISA) decomposed the global bivariate Moran’s I into pixel-level cluster assignments, enabling spatially explicit identification of concordant and discordant zones (Fig. 8; Table 11).

**Fig. 8.**
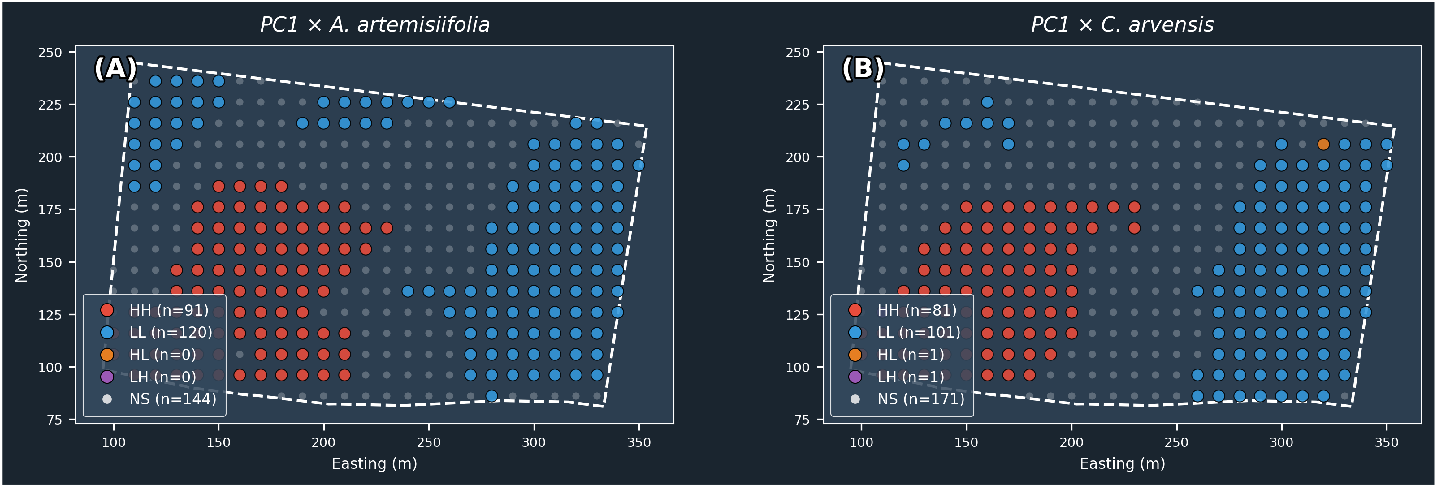
Bivariate LISA cluster maps (seedling phase, Santa Rosa paddock). (A) *A. artemisiifolia*: HH = 109, LL = 126, HL = 22, LH = 1; 258 of 380 pixels (67.9%) significant. (B) *L. culinaris*: HH = 92, LL = 99, HL = 28, LH = 0; 219 of 380 pixels (57.6%) significant. HH (high-high) clusters indicate priority treatment areas where both satellite signal and weed density are elevated; LL (low-low) clusters are confirmed clean zones. Spatial weights: DistanceBand (15 m threshold), row-standardised; 999 conditional permutations, *p <* 0.05.

**Table 11.**
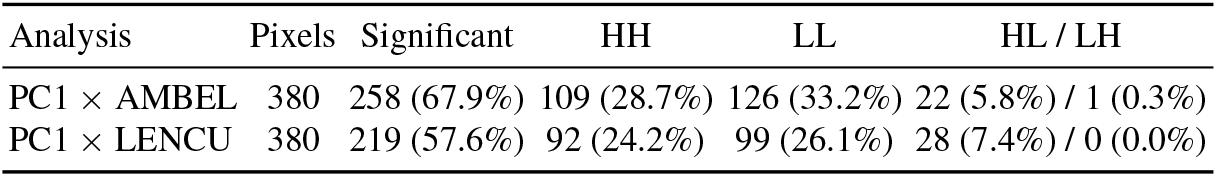
Bivariate LISA cluster analysis of PRESTO PC1 versus kriged weed density. HH = High-High (hot spot); LL = Low-Low (cold spot); HL = High-Low (spectral outlier); LH = Low-High (missed infestation). Significance assessed at *α* = 0.05 with 999 permutations.

For AMBEL, 67.9% of all paddock pixels were classified as statistically significant bivariate clusters, of which 28.7% fell in the High-High (HH) quadrant and 33.2% in the Low-Low (LL) quadrant. HH pixels are zones where the satellite embedding signal and ground-truth weed density jointly indicate heavy infestation; LL pixels are zones where both data sources agree on low weed pressure. The remaining significant pixels comprised a modest population of spectral outliers (HL: 5.8%), where the embedding value was elevated but weed density remained low, and a single Low-High (LH) pixel (0.3%), where weed density was high but the satellite signal did not flag it.

The LENCU analysis exhibited a similar pattern, with 57.6% significant pixels, 24.2% HH, 26.1% LL, 7.4% HL, and zero LH pixels. The near-total absence of LH pixels in both analyses indicates that the satellite-derived embedding system exhibited virtually no blind spots—it almost never failed to flag pixels where ground-truth weed density was elevated. The HL (false alarm) rate was 6–7%.

#### Cross-variograms

Cross-variograms were computed to quantify the spatial scale at which the embedding-weed co-variation reaches its maximum extent (Fig. 9). The PC1 auto-variogram stabilized at a range of approximately 84 m. The cross-variograms between PC1 and kriged weed density yielded ranges of approximately 90 m for both AMBEL and LENCU. The close agreement between the auto-variogram and cross-variogram ranges indicates that the satellite embedding field and the weed density field share the same characteristic spatial scale of variation.

**Fig. 9.**
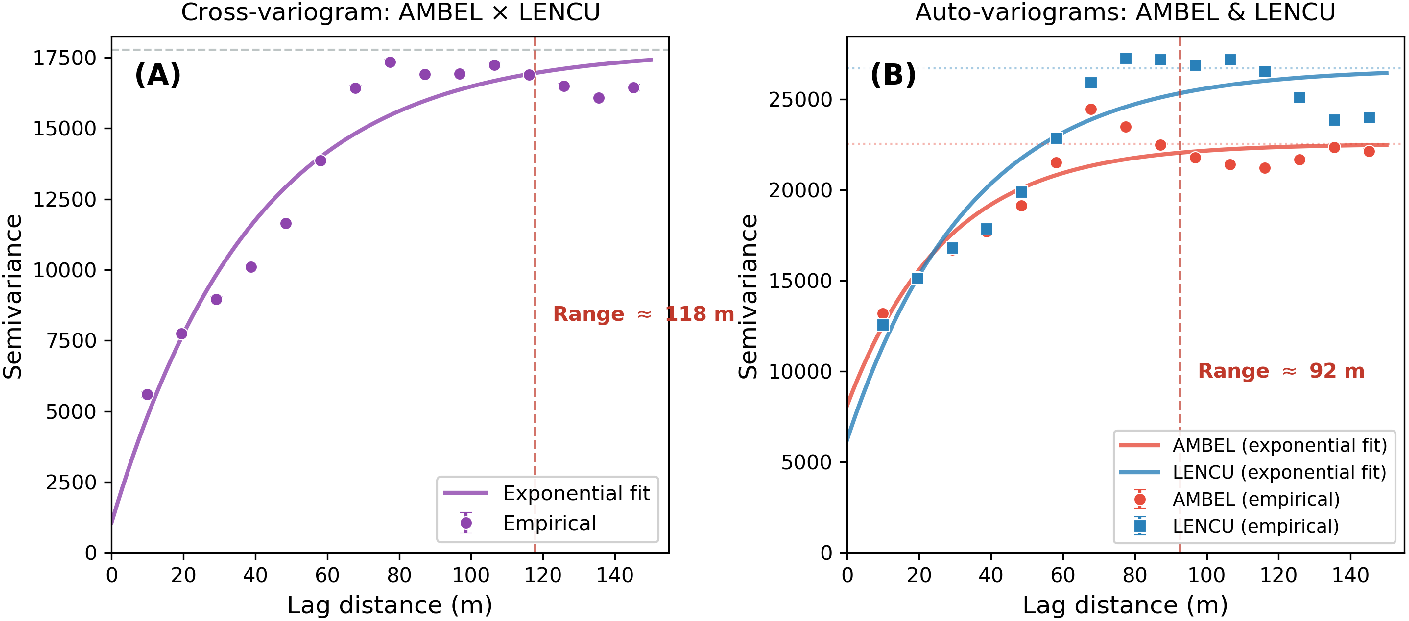
Cross-variograms (seedling phase). (A) PC1 vs. *A. artemisiifolia* and *L. culinaris*, and (B) auto-variograms. Exponential model fitted to empirical semivariance at 15 lag bins. Practical range *≈* 90 m for cross-variograms; PC1 auto-variogram range *≈* 84 m. Dashed vertical line marks the 90 m range. The spatial grain of the satellite-weed relationship indicates that the spectral bridge operates at management-relevant scales.

The 90 m range defines the natural patch size of the satellite-weed relationship within this paddock. The operational implications of this spatial scale are discussed in Section 5.3.

#### Geographically weighted regression

Geographically Weighted Regression (GWR) (Fotheringham et al., 2002) was applied with PC1, PC2, and PC3 as predictors and kriged weed density as the dependent variable to test for spatial non-stationarity in the embedding-density relationship (Table 12; Fig. 10).

**Fig. 10.**
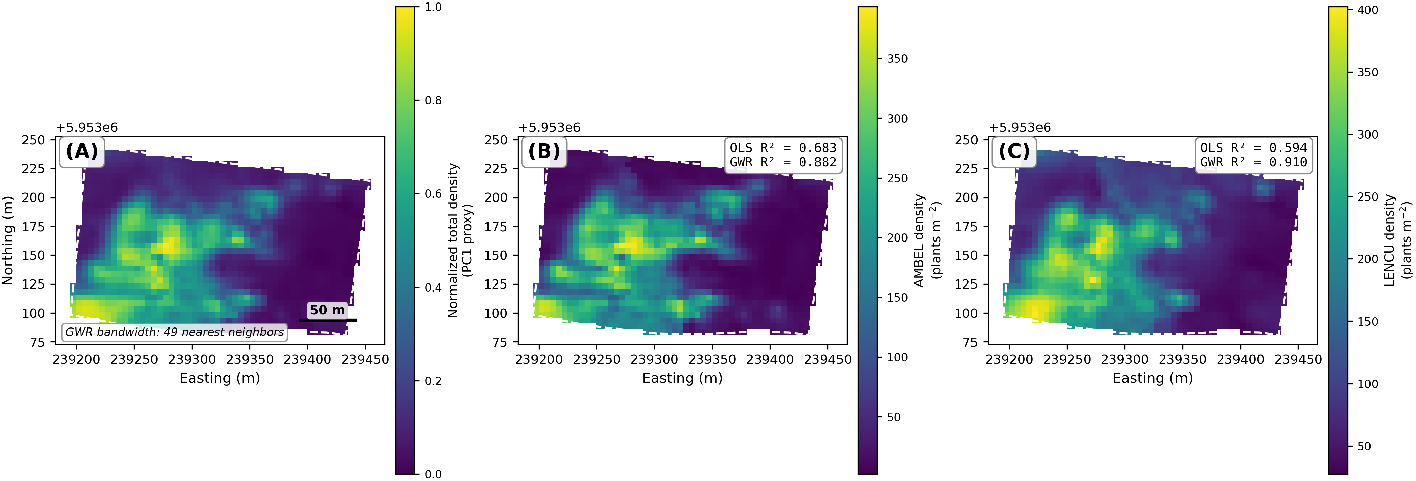
Side-by-side spatial comparison (seedling phase) in the Santa Rosa lentil paddock. (A) Presto PC1, (B) *A. artemisiifolia* kriging density, (C) *L. culinaris* kriging density. GWR with adaptive band-width (49 nearest neighbours) improved fit over global OLS: *R*^2^ = 0.683 *→* 0.882 for *A. artemisiifolia* and *R*^2^ = 0.594 *→* 0.910 for *L. culinaris*. Viridis colourmap; dashed white line indicates paddock boundary. Spatial correspondence between satellite embeddings and ground-truth weed density confirms the spectral bridge.

**Table 12.**
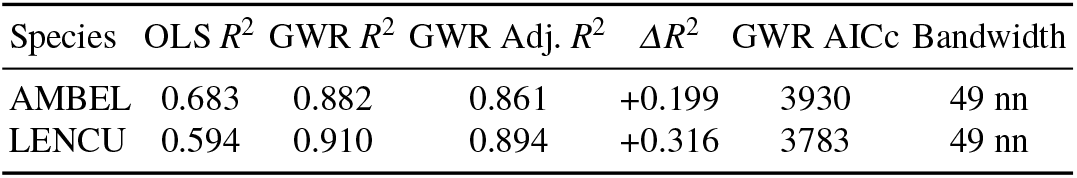
Geographically Weighted Regression (GWR) versus global Ordinary Least Squares (OLS) for predicting kriged weed density from PRESTO principal components. AICc = corrected Akaike Information Criterion; nn = nearest neighbors.

| Species | OLS $R^2$ | GWR $R^2$ | GWR Adj. $R^2$ | $\Delta R^2$ | GWR AICc | Bandwidth |
| --- | --- | --- | --- | --- | --- | --- |
| AMBEL | 0.683 | 0.882 | 0.861 | +0.199 | 3930 | 49 nn |
| LENCU | 0.594 | 0.910 | 0.894 | +0.316 | 3783 | 49 nn |

GWR dramatically outperformed the global OLS baseline for both species. For AMBEL, the *R*^2^ increased from 0.683 under OLS to 0.882 under GWR, a gain of +0.199; for LENCU, the improvement was even larger, from 0.594 to 0.910 (+0.316). The adjusted *R*^2^ values (0.861 and 0.894, respectively) confirmed that the improvement was not an artifact of model complexity. AICc values of 3930 (AMBEL) and 3783 (LENCU) were substantially lower than those of the corresponding OLS models, providing independent confirmation that the local model achieved a genuine improvement in fit. The adaptive bandwidth of 49 nearest neighbors indicates that the regression coefficients vary over distances consistent with the 90 m cross-variogram range, reinforcing the geostatistical evidence for patch-scale spatial non-stationarity in the embedding-weed relationship.

The implications of this 20–32 percentage-point improvement are discussed in Section 5.3.

## 5 Discussion

### 5.1 Automated Surveillance at Three Spatial Scales

The primary objective of this chapter was to develop surveillance tools for *A. artemisiifolia* that overcome the landscape-scale detection bottleneck described in the Introduction. The results presented in Section 4 demonstrate that each analytical layer— field-scale detection, cross-domain generalization, and satellite-scale integration— addresses a distinct segment of the surveillance challenge, yet none constitutes a self-sufficient solution under variable field conditions. The YOLOv11l detector achieved mAP_50_ = 0.886 on its training domain (Table 6), confirming that deep learning can reliably locate ragweed and associated broadleaf weeds in ground-level imagery, but its near-complete failure on international data (mAP_50_ = 0.108; Table 8) revealed that detection performance is tightly coupled to acquisition context. Multi-domain training recovered international performance to 0.874 at the cost of a 14.4 percentage-point regression on the Chilean test set, illustrating the well-known trade-off between domain breadth and domain specificity. At the satellite scale, geographically weighted regression explained 88–91% of the local variance in weed density from PRESTO embeddings alone (Table 12), yet this explanatory power was demonstrated on a single 3.42 ha paddock and cannot yet be assumed to transfer to other sites, soil types, or crop rotations.

The training progression itself yielded instructive patterns for practitioners. The comparison between Runs 2 and 3 demonstrates that, holding the training set constant, larger batches provide more stable gradient estimates per optimization step, yielding both faster convergence and better final metrics. Run 3’s batch size of 64 on dual GPUs completed training in 39.9 min while surpassing Run 2 (batch size 8, 164.0 min) by 1.7 points in mAP_50_ and 5.1 points in mAP_50-95_—a result that argues for maximizing effective batch size before investing in higher resolution or longer schedules. Run 4’s degradation trajectory, by contrast, illustrates the risk of catastrophic forgetting: the default learning rate of 0.01 proved sufficiently aggressive to overwrite the feature representations learned during Run 3 pre-training, producing immediate performance collapse from epoch 1 onward. This pattern is characteristic of transfer learning scenarios in which the initial learning rate exceeds the basin of attraction of the pre-trained solution. Perhaps most instructive for operational deployment, Run 4 detected 58% more ragweed plants in the Santa Rosa field images than Run 3 despite its lower aggregate validation metrics (mAP_50_ of 0.839 versus 0.886). This discrepancy between validation-set performance and site-specific detection counts underscores that practitioners cannot rely exclusively on held-out metrics when selecting models for deployment in specific field contexts; targeted evaluation on representative imagery from the intended deployment environment remains essential.

The deliberate framing of these components as a *toolkit* rather than a finished system reflects both an epistemological and a practical stance. From an epistemological perspective, each method was explored, tested, and documented with its performance boundaries explicitly quantified so that future users can judge applicability to their own conditions. From a practical perspective, the value of the work lies in the *integration* of detection, embedding-based domain diagnostics, and spatially explicit satellite prediction into a reproducible open-source pipeline. The package source code, including its geoprocessing modules, is released as a community deliverable (operational scripts and trained weights are available from the corresponding author upon request), enabling other research groups to adapt, extend, or replace individual modules as climate variability reshapes weed communities and management requirements across geographies.

### 5.2 The Spectral Bridge Is Confirmed

The central finding of the satellite-scale analysis is that foundation model embeddings derived from six months of multi-sensor observations at 10 m resolution can predict sub-field weed density with quantitative precision. This result establishes what we term a *spectral bridge*: a measurable, spatially coherent link between the field-scale detection layer (individual plant counts from YOLO+SAHI at centimeter resolution) and the landscape-scale monitoring layer (PRESTO embeddings from Sentinel-2 at decameter resolution).

The mechanism underlying this bridge is direct spectral contribution. Above-ground broadleaf species—both the weed *Ambrosia artemisiifolia* and the host crop *Lens culinaris*—add green biomass within each 10 m pixel, elevating the vegetation signal in infested zones relative to clean crop areas. Single-date NDVI captured this effect most transparently, achieving the highest point-wise correlations (*r* = 0.890 for LENCU; *r* = 0.837 for AMBEL; Table 10). That a single spectral index from a single acquisition outperformed PRESTO PC1 (*r* = 0.717 and 0.739) in point-wise terms is informative: when the analyst can select the acquisition date to coincide with peak spectral contrast, simpler approaches remain competitive. This contrasts with root-parasitic species such as *Orobanche* spp., where the satellite signal is mediated by indirect crop stress responses rather than the weed’s own canopy, and where single-date indices are unlikely to achieve comparable correlations.

The POLAV (*Polygonum aviculare*) result delimits the applicability of this spectral bridge. With *r* = 0.194 (95% CI [0.095, 0.289]), POLAV failed to establish a meaningful satellite–weed association despite being detected in the same imagery and processed through the identical analytical pipeline. The mechanism is biomass visibility: *P. aviculare* is a prostrate, low-biomass species whose spectral contribution is overwhelmed by the combined crop and dominant weed signal within each 10 m pixel. This species-specific failure carries a direct management consequence: satellite-based weed mapping cannot replace ground-level scouting for species that contribute minimal above-ground biomass. The spectral bridge, as demonstrated here, is applicable to upright, biomass-dominant broadleaf species whose canopy presence measurably alters the pixel-level vegetation signal.

However, the comparison between NDVI and PRESTO is not symmetric. Single-date indices are vulnerable to cloud contamination, phenological timing errors, and the assumption—difficult to verify *a priori*—that the chosen date corresponds to peak weed–crop spectral contrast. PRESTO, by encoding the full six-month temporal trajectory, is inherently resilient to individual suboptimal dates (Tseng et al., 2023). Moreover, the temporal embeddings carry spatial structure that single indices cannot: bivariate Moran’s *I* = 0.706 (*p* = 0.001) for the PC1–AMBEL association confirmed that the satellite–weed relationship is spatially coherent across the paddock, not merely an aggregate statistical artifact (Anselin, 1995). The LISA decomposition reinforced this interpretation: 258 of 380 pixels (67.9%) were classified into significant bivariate clusters, with nearly zero Low-High pixels (Table 11), indicating that the satellite system exhibited virtually no blind spots where ground-truth infestations went undetected. To our knowledge, this represents the first demonstration that temporal foundation model embeddings from Sentinel-2 can predict broadleaf weed density at sub-field resolution (5 m).

### 5.3 Spatial Non-Stationarity Matters

The most striking quantitative result of the satellite-scale analysis was the magnitude of the improvement achieved by allowing regression coefficients to vary geographically. Geographically weighted regression (Fotheringham et al., 2002) increased explained variance from *R*^2^ = 0.683 under global OLS to *R*^2^ = 0.882 under GWR for AMBEL (+0.199), and from *R*^2^ = 0.594 to *R*^2^ = 0.910 for LENCU (+0.316; Table 12). These gains, amounting to 20–32 percentage points, were confirmed by substantially lower AICc values and by adjusted *R*^2^ estimates (0.861 and 0.894) that rule out overfitting as an explanation. The message is unambiguous: the relationship between satellite embeddings and weed density is not spatially stationary, and any operational system that assumes a single global regression will systematically underestimate its own predictive capacity.

Several drivers of this non-stationarity are plausible within the Santa Rosa paddock. Soil heterogeneity—including contrasting soil texture and mineralogy zones characteristic of alluvial soils in central Chile—creates spatially varying growing conditions that modulate both crop vigor and weed competitive success. Microclimate gradients associated with topography, fence lines, and irrigation infrastructure further diversify the embedding–weed relationship across the 3.42 ha extent. Management history, including differential herbicide application and tillage patterns, introduces abrupt spatial transitions in weed pressure that cannot be captured by a single set of regression coefficients.

The adaptive bandwidth of 49 nearest neighbors selected by GWR is consistent with the approximately 90 m cross-variogram range that defined the characteristic patch size of the satellite–weed co-variation (Section 4.3). Both statistics converge on the same interpretation: the natural spatial grain of the weed infestation system in this paddock operates at scales of roughly 80–100 m. Management zones smaller than this range would sub-resolve the underlying spatial structure, while zones larger than it would mask heterogeneity that the data are capable of capturing.

The implications extend beyond this single paddock. Under the non-stationary climatic conditions that frame this volume, spatial heterogeneity in agricultural systems is expected to intensify as temperature and precipitation regimes become more variable (Lake et al., 2017). If the embedding–weed relationship is already markedly nonstationary within a uniform 3.42 ha lentil field, then larger and more heterogeneous production units—the typical operational context for precision weed surveillance—will require spatially adaptive models as a prerequisite, not an optional refinement. Global regressions, however convenient, are best regarded as diagnostic baselines rather than operational endpoints.

### 5.4 Domain Shift as the Primary Deployment Risk

The cross-domain experiment delivered a clear and operationally consequential message: the primary obstacle to deploying weed detection models in new geographies is not species recognition but acquisition conditions. Camera hardware, illumination regime, background vegetation, and perspective angle drove the dominant axes of variation in the ResNet50 embedding space (Section 4.2), while species morphology played a secondary role. When the Chilean-only model was applied to international imagery without adaptation, mAP_50_ collapsed to 0.108—not through gradual degradation proportional to visual novelty, but through a categorical failure to produce any detections at all. This result extends the Chile–Denmark domain gap documented by León et al. (2024b) from a bilateral comparison to a nine-database, four-country quantification, confirming that the phenomenon is systematic rather than site-specific.

The embedding-based deployment-gate framework proposed in Section 3.2.1 was empirically validated by these results. All Chilean-to-international MMD values fell in the 0.27–0.43 range, which the gate framework classified as requiring augmentation or active learning before deployment (Gretton et al., 2012). Phase 1 confirmed the prediction of failure; Phase 2 confirmed the prescribed remedy. The practical implication is direct: before deploying a trained model in a new geography, practitioners should compute pairwise MMD between the training distribution and a small unlabeled sample from the target environment—a procedure requiring no annotations, only images and a frozen feature extractor. Values below 0.15 permit direct deployment; values above 0.30 should trigger data collection from the target domain.

Multi-domain training recovered international performance to mAP_50_= 0.874 (+76.6 percentage points), but incurred a 14.4 percentage point regression on the Chilean test set (0.886 to 0.742). This trade-off between generality and specialization is not unique to weed detection; it is a well-characterized property of domain generalization in computer vision. However, the magnitude of the Chilean regression exceeded the pre-registered five-point guard-rail, indicating that naive pooling of domains is insufficient. Curriculum-based or domain-weighted sampling strategies, combined with extended training schedules, represent the most promising avenues for recovering domain-specific precision without sacrificing cross-domain breadth.

### 5.5 Species Discrimination at Altitude

The class inversion observed between ground-level and drone-altitude detection constitutes, to our knowledge, a novel finding that has not been previously reported in the weed detection literature. The original ground-trained model assigned 11.6% of drone detections to AMBEL and 65.4% to LENCU; after retraining with drone-specific annotations, these proportions reversed to 44.9% and 28.1%, respectively (Table 7). Yet the combined broadleaf detection rate—the sum of AMBEL and LENCU—remained stable at 73–84% across all three models evaluated. This pattern reveals that the models reliably detect broadleaf objects at drone altitude but redistribute class labels depending on training context.

The mechanism is morphological convergence at altitude. At ground level, *Ambrosia artemisiifolia* and *Lens culinaris* are distinguishable by leaf architecture and growth habit. At 20–30 m altitude, these diagnostic features collapse below pixel resolution, and both species present as small, irregularly shaped green patches against the soil background. The model is therefore solving a density estimation problem rather than a species identification problem when operating on drone imagery.

This finding has two practical consequences. First, drone-based detection should be reported at the broadleaf-aggregate level rather than at the species level, because species assignments at altitude are unreliable and potentially misleading for management decisions such as herbicide selection. Second, a two-stage detection strategy is warranted: satellite screening identifies hot-spot zones across the paddock, drone surveys quantify weed density within those zones at high spatial resolution, and targeted ground verification confirms species identity where it matters for treatment decisions. This hierarchical approach allocates the most expensive observation—ground-level expert identification—only to locations where species-level discrimination is both necessary and achievable.

Under climate change, the relevance of species-level identification will intensify. As weed communities shift poleward and novel species establish in previously unaffected regions, management strategies will increasingly depend on knowing *which* species is present, not merely that weeds are present. The two-stage strategy positions drone density mapping as a rapid triage tool while preserving species-level resolution where it can actually be resolved.

### 5.6 Climate Variability Implications

The tools developed in this chapter were designed for a specific weed species in a specific cropping system, but the underlying challenges—domain shift, spatial non-stationarity, and scale bridging—are generic consequences of working in non-stationary environments. Climate change amplifies each of these challenges, and the toolkit architecture responds to them directly.

#### Range expansion

Rising temperatures and shifting precipitation patterns are extending the geographic range of *A. artemisiifolia* into previously uncolonized regions across Europe, South America, and East Asia (Lake et al., 2017). Long-term data from the Broadbalk experiment confirm that weed competitive advantage over crops has increased since 1969 in response to warming (Storkey et al., 2021), and ecological niche models show that ragweed’s niche expansion in South America (expansion index = 0.407) exceeds that on any other continent (Song et al., 2023). Each new colonization front represents a domain shift event: local vegetation backgrounds, illumination conditions, and associated crop types differ from those in the model’s training distribution. The cross-domain experiment demonstrated that domain diversity in the training pool—not dataset size—is the decisive factor for surviving such shifts (Section 4.2). The deployment-gate framework provides a quantitative mechanism for deciding when a model can be transferred to a new region and when retraining is required.

#### Phenological shifts

Elevated CO_2_ concentrations and warmer temperatures alter ragweed phenology, extending the growing season and increasing pollen production per plant (Ziska et al., 2003). These effects are already measurable: pollen seasons in North America have lengthened by approximately 20 days since 1990 (Anderegg et al., 2021), and projections indicate a further 16–40% increase from climate alone—or up to 200% when CO_2_ fertilization is included (Zhang & Steiner, 2022). PRESTO embeddings encode six-month spectral-temporal trajectories rather than single-date snapshots (Tseng et al., 2023), making them inherently sensitive to phenological timing. As growing seasons shift, the temporal signature captured by the foundation model will track these changes without requiring manual recalibration of acquisition dates.

#### Non-stationary management

The GWR results (Section 4.3) demonstrated that the relationship between satellite-derived embeddings and weed density varies spatially across a single 3.42 ha paddock, with global models underestimating predictive power by 20–32 percentage points. Climate variability will amplify this spatial heterogeneity by introducing micro-climatic gradients, differential soil moisture responses, and site-specific phenological offsets. Spatially varying models are therefore not a statistical refinement but a practical necessity for operational weed surveillance under changing conditions.

#### Public health

Common ragweed is the primary source of allergenic pollen in late summer across temperate regions, and pollen loads are projected to double under moderate warming scenarios by 2050 (Lake et al., 2017), with associated healthcare costs exceeding EUR 7 billion annually in Europe alone (Schaffner et al., 2020). Beyond health impacts, ragweed causes yield losses of up to 83.7% in soybean (Hall et al., 2021), and approximately 90% of pollen grains deposit within 100 m of the source (Katz & Batterman, 2019), reinforcing the case for field-level source reduction. Proactive surveillance of agricultural ragweed populations before the pollen season— using the LISA hot-spot clusters (HH zones) identified in Section 4.3 as priority spray targets—represents a source-reduction strategy that addresses the problem at its origin rather than managing symptoms downstream.

#### Converging evidence for AI-assisted monitoring

Climate warming is also eroding classical biocontrol options: thermal stress reduces the efficacy of *Ophraella* leaf beetles released against ragweed through genetic and metabolomic changes in the host plant (Sun et al., 2022), strengthening the case for detection-based management. At the landscape scale, satellite classification combined with aerobiological data has been used to map ragweed habitat in Serbia (Lugonja et al., 2019), and machine learning pollen forecasting models have identified the absence of spatially explicit “source maps” as a key bottleneck for operational prediction (Zewdie et al., 2019a;b). Recent work linking satellite-derived aerosol properties to continental-scale pollen mapping (Zhang et al., 2026) further underscores the operational value of the georeferenced density surfaces produced by the present toolkit. Complementary work in precision agriculture robotics underscores the generality of the domain shift challenge: transfer learning from citizen science photographs to UAV species identification (Soltani et al., 2022), domain generalization for crop–weed segmentation across fields (Weyler et al., 2023), and unsupervised domain adaptation for field-to-field weed mapping (Magistri et al., 2023) all confirm that bridging the gap between training and deployment environments is the central obstacle to scalable weed monitoring—the same obstacle addressed by the embedding-based deployment gate developed here.

#### Open toolkit

The complete pipeline—from YOLO training configurations to SAHI inference scripts, embedding extraction, and spatial analysis workflows—is released as an open repository. This design decision reflects the conviction that tools for managing non-stationary biological systems must themselves be adaptable. As conditions change across seasons, sites, and years, the community can retrain models, recalibrate deployment gates, and extend the spatial analysis to new paddocks without rebuilding the pipeline from scratch.

### 5.7 Limitations and Future Work

Several limitations constrain the generalizability of the present results and define directions for future research.

#### Anomaly confounding

The satellite analysis assumes that spectral deviations from the crop baseline are attributable to weed presence. In practice, nutrient deficiency, waterlogging, disease, and soil variability can produce similar spectral anomalies. The PRESTO embeddings do not discriminate between these sources of within-field heterogeneity, and the high GWR *R*^2^ values (0.882–0.910) may partly reflect spatial confounding between weed density and underlying soil or management gradients rather than a direct causal link between the spectral signal and weed biomass. Resolving this ambiguity will require multi-temporal analysis at finer temporal resolution, potentially leveraging the self-attention layers within the PRESTO architecture to identify which spectral bands and time steps carry the weed-specific signal.

#### Single-site satellite validation

All satellite-scale results are derived from a single 3.42 ha paddock (Santa Rosa) during one growing season. While the within-paddock analysis is robust—380 spatially aligned pixels with ground-truth kriging from 1,685 drone photograms—the external validity of the GWR coefficients, LISA cluster patterns, and cross-variogram ranges has not been tested. Multi-site, multi-season replication is essential before the spectral bridge can be considered operational.

#### Chilean regression

The 14.4 percentage point decline in Chilean mAP_50_ under combined training exceeds acceptable tolerances for operational deployment. The regression is partly attributable to training constraints (50 vs. 100 epochs, single vs. dual GPU) rather than fundamental incompatibility, but domain-balanced sampling and longer training schedules must be validated before the combined model can replace the domain-specific one in Chilean production systems.

#### Orthomosaic pipeline

The orthomosaic detection workflow reached only the tile extraction and MMD assessment stages (phases 0–3 of 8). The 2.4 GB mosaic (73,000 *×* 41,000 px, 2,151 tiles) and the 80 tiles selected for active learning represent a methodological proof of concept, but full annotation, retraining, and paddock-scale inference remain incomplete.

#### Detector ground truth

All downstream analyses—kriging density surfaces, LISA clusters, spectral–weed correlations, GWR models—treat YOLOv11 detection counts as unchallenged ground truth. Detection errors, both false positives and false negatives, propagate into all spatial statistics and may inflate or deflate the reported correlations and explained variance. A manual validation subsample of 50–100 randomly selected photograms, scored by an independent observer, would quantify this error floor and enable formal uncertainty propagation through the analysis chain.

#### Active learning

The deployment-gate framework identifies when retraining is needed but does not yet close the loop by automatically selecting, annotating, and incorporating target-domain samples. Integrating uncertainty-based active learning with the MMD gate would create a self-improving system that requests human annotation only for the most informative images.

#### Future directions

Three extensions are prioritized. First, exploiting PRESTO’s temporal self-attention mechanism to disentangle weed, disease, and nutrient stress signals within the same paddock. Second, expanding the detection pipeline to additional cropping systems—wheat, fallow, maize, and tomato datasets already exist in the repository—to test whether the toolkit generalizes beyond the lentil–ragweed system. Third, implementing real-time SAHI inference on drone video feeds to enable on-the-fly density mapping during survey flights, eliminating the current post-processing delay between data acquisition and management prescription.

## 6 Conclusions

This chapter presented an integrated surveillance toolkit for detecting, mapping, and characterizing *Ambrosia artemisiifolia* infestations across spatial scales ranging from individual plants to satellite pixels. Five principal findings emerged:

1. **First ragweed detection model, field-scale baseline.** To our knowledge, this is the first deep learning detection model for common ragweed. YOLOv11l trained on 4,335 ground-level images (adult phase) with SAHI sliced inference at 640 px achieved mAP_50_ = 0.886, mAP_50-95_ = 0.744, precision of 0.934, and recall of 0.808 (Table 6). This establishes a quantitative baseline for automated ragweed detection and enables the cross-domain and satellite-scale analyses that follow.
2. **Domain shift is predictable and recoverable.** A Chilean-only model collapsed to mAP_50_ = 0.108 on international imagery (adult phase)—a categorical failure, not gradual degradation. The embedding-based deployment-gate framework (MMD thresholds: *<*0.15 deploy, 0.15–0.30 augment, 0.30–0.45 active learn, *>*0.45 retrain) correctly predicted this failure. Multi-domain training recovered international performance to 0.874, demonstrating that domain diversity, not dataset size, drives cross-domain generalization (Table 8). This framework enables practitioners to assess model transferability *before* deployment in new geographies.
3. **Satellite embeddings predict weed density; spectral bridge confirmed.** PRESTO temporal embeddings (seedling phase), analyzed through geographically weighted regression, explained 88.2% (AMBEL) and 91.0% (LENCU) of the local variance in kriged weed density (Table 12). This demonstrates that sub-field density information derived from drone imagery can be successfully linked to 10 m satellite pixels through foundation model embeddings and spatially varying regression, enabling paddock-level surveillance from remote sensing data alone. Bivariate LISA analysis confirmed spatially coherent hot-spot concordance, with near-zero blind spots (LH pixels: 0–0.3%).
4. **Spatial non-stationarity is essential, not optional.** Global regression models underestimated the predictive power of satellite embeddings by 20–32 percentage points relative to GWR (Table 12). Cross-variogram analysis identified 90 m as the characteristic spatial scale of the embedding-weed relationship, defining the minimum meaningful surveillance zone size. Operational systems assuming spatially uniform relationships will systematically underestimate their own capacity for spatial discrimination.
5. **Open surveillance pipeline and LISA-guided source reduction.** The complete pipeline—training configurations, SAHI inference scripts, embedding extraction workflows, and spatial analysis code—is released as an open repository (trained model weights are available from the corresponding author upon request), enabling adaptation to new species and geographies. The bivariate LISA High-High clusters identify spatial foci of *A. artemisiifolia* infestation at sub-field resolution, enabling targeted source-reduction interventions before reproductive maturity and peak pollen dispersal. Given that ragweed pollen sensitization affects over 50 million Europeans and pollen loads are projected to double under moderate warming (Lake et al., 2017), field-level source maps that prioritize surveillance and treatment targets represent a direct contribution to public health strategies addressing the problem at its agricultural origin.

Taken together, these findings demonstrate that automated surveillance for invasive weeds can span from individual plants to satellite-scale observations through the integration of deep learning detection, foundation model embeddings, and spatially explicit geostatistics. The urgency of this integration is underscored by projections of a 16–200% increase in allergenic pollen loads by the end of the century (Zhang & Steiner, 2022) and by evidence that weeds adapt faster than crops to changing climatic conditions (Anwar et al., 2021). The surveillance system is imperfect—the 14.4 percentage-point Chilean regression under combined training, the POLAV failure revealing biomass thresholds for satellite detection, and the incomplete orthomosaic pipeline all indicate that further development is required. Yet the core capabilities— automated field-scale plant detection, domain-gap prediction, and satellite-scale density estimation—have been demonstrated and validated.

The modular design is not a convenience but a necessity. Under the non-stationary conditions that climate change creates—shifting species ranges, altered phenology, amplified spatial heterogeneity—surveillance systems designed for fixed conditions will degrade as environments change. Long-term evidence confirms this trajectory: weed competitive advantage over crops has increased steadily since the late 1960s (Storkey et al., 2021). The toolkit paradigm enables adaptation: practitioners and researchers can retrain models with new annotated data, recalibrate deployment gates as new geographies reveal different domain gaps, and extend spatial analyses to different paddocks without rebuilding the pipeline from scratch.

## Acknowledgements

The authors thank Prof. Noureddine Benkeblia for the editorial invitation to contribute to this volume. Field data collection and annotation were supported by Sebastián Medina and Alberto Espinoza at the Santa Rosa site. Computational resources were provided by the Instituto de Investigaciones Agropecuarias (INIA), Chillán, Chile. The international ragweed datasets were made available through their respective open-access repositories: the authors of the ND Individual, Michigan 3-Season, WeedCrop PrecAg, ND Aerial, Purdue 4Weed, and WeedCube USDA datasets are gratefully acknowledged. Satellite imagery was accessed via the Copernicus Data Space Ecosystem. The PRESTO foundation model was developed by the WorldCereal consortium and made available under open license.

The authors used Claude (Anthropic) as a large language model assistant for manuscript drafting, code development, and data analysis pipeline construction. All scientific content, experimental design, data collection, analysis, and interpretation were performed by the authors, who take full responsibility for the accuracy and integrity of the work. All figures are data-driven visualizations generated from analytical scripts; no figures were created using generative AI. This work was supported by the Ministry of Agriculture of Chile through the Fundación para la Innovación Agraria (FIA).

## Ethics Approval

Not applicable. This study did not involve human participants or animals.

## Data Availability

The weed detection datasets used in this study are available from their respective open-access repositories as cited in Section 2.2. The Chilean field imagery is available from the corresponding author upon reasonable request.

## Code Availability

The package source code, analysis workflows, and documentation are released as an open-source Python package (ragweed-ai-toolkit, MIT license) comprising eight analytical modules and eight command-line entry points: https://github.com/agroia-lab/ragweed-ai-toolkit. The 27 operational scripts and the trained model weights are available from the corresponding author upon reasonable request.

## About the Author

**Lorenzo F. León Gutiérrez** is Weed Science and Technology Program Director at the Centro Regional de Investigación (CRI) INIA Quilamapu, Instituto de Investigaciones Agropecuarias, Chillán, Chile. An agronomist with a Magister (Mg.Sc.) degree and over twenty years of experience at INIA (2006–present), his career has progressed from precision farming, spatial variability characterisation, and non-destructive spectroscopic analysis of agricultural products (vis/NIR, fluorescence) to his current focus on deep learning, geospatial statistics, and remote sensing for integrated weed management under climate variability. He directs research on AI-driven workflows for weed detection and mapping in cereal, legume, and row-crop systems, spanning object detection in field and aerial imagery (YOLOv11, SAHI), embedding-based cross-domain transfer learning across geographies, and multi-scale spatial analysis linking drone-level detections to satellite-derived indicators through foundation models. His current work centres on *Ambrosia artemisiifolia* (common ragweed) as a climate-change marker species, bridging agricultural management with public health outcomes through scalable surveillance frameworks. His work is guided by the principle that effective agricultural AI must be open-source, reproducible, and accessible to practitioners and researchers across diverse production systems.

## List of Abbreviations

AICc: Corrected Akaike Information Criterion
AMBEL: EPPO code for *Ambrosia artemisiifolia* (common ragweed)
AMP: Automatic Mixed Precision
BSI: Bare Soil Index
CIoU: Complete Intersection over Union
CLI: Command-Line Interface
COCO: Common Objects in Context (benchmark dataset)
DFL: Distribution Focal Loss
EPPO: European and Mediterranean Plant Protection Organization
EVI: Enhanced Vegetation Index
EXIF: Exchangeable Image File Format
GNDVI: Green Normalized Difference Vegetation Index
GPS: Global Positioning System
GSD: Ground Sampling Distance
GWR: Geographically Weighted Regression
HH: High-High (LISA cluster concordant hot spot)
HL: High-Low (LISA cluster spatial outlier)
IoU: Intersection over Union
LENCU: EPPO code for *Lens culinaris* (lentil)
LH: Low-High (LISA cluster spatial outlier)
LISA: Local Indicators of Spatial Association
LL: Low-Low (LISA cluster concordant cold spot)
mAP: mean Average Precision
MMD: Maximum Mean Discrepancy
NDMI: Normalized Difference Moisture Index
NDVI: Normalized Difference Vegetation Index
NDWI: Normalized Difference Water Index NMS Non-Maximum Suppression
OLS: Ordinary Least Squares
PCA: Principal Component Analysis
POLAV: EPPO code for *Polygonum aviculare* (prostrate knotweed)
POLPE: EPPO code for *Polygonum persicaria* (pale persicaria)
PRESTO: Pretrained Remote Sensing Transformer
SAHI: Slicing Aided Hyper Inference
SAVI: Soil-Adjusted Vegetation Index
SGD: Stochastic Gradient Descent
SWIRd: Shortwave Infrared Difference Index
t-SNE: t-Distributed Stochastic Neighbor Embedding
UAV: Unmanned Aerial Vehicle
UMAP: Uniform Manifold Approximation and Projection
YOLO: You Only Look Once (object detection architecture)

